# LLAMA: a robust and scalable machine learning pipeline for analysis of cell surface projections in large scale 4D microscopy data

**DOI:** 10.1101/2020.12.10.420414

**Authors:** James G. Lefevre, Yvette W. H. Koh, Adam A. Wall, Nicholas D. Condon, Jennifer L. Stow, Nicholas A. Hamilton

## Abstract

We present LLAMA, a pipeline for systematic analysis of terabyte scale 4D microscopy datasets. Analysis of individual biological structures in imaging at this scale requires efficient and robust methods which do not require human micromanagement or editing of outputs. To meet this challenge, we use a machine learning method for semantic segmentation, followed by a robust and configurable object separation and tracking algorithm, and the generation of detailed object level statistics. Advanced visualisation is a key element of LLAMA: we provide a specialised software tool which supports quality control and optimisation as well as visualisation of outputs. LLAMA was used in a quantitative analysis of macrophage surface membrane projections (filopodia, ruffles, tent-pole ruffles) examining the differential effects of two interventions: lipopolysaccharide (LPS) and macrophage colony stimulating factor (CSF-1). Distinct patterns of increased activity were identified. In addition, a continuity of behaviour was found between tent pole ruffling and wave-like ruffling, further defining the role of filopodia in ruffling.

## Introduction

In vertebrate cells the cell surface can be dynamically deformed to produce a variety of membrane projections that are used for interactions with other cells and the microenvironment. As innate immune cells, macrophages are well known for their ability to extend dramatic filopodia, ruffles and phagocytic cups which contribute to their roles in immune surveillance and defence (Stow & Condon, 2016). Continuous, dynamic ruffling is a feature of the macrophage surface, occurring constitutively, it is further enhanced by cell activation (Condon et al., 2018). Large membrane ruffles can close, entrapping and internalising fluid-phase cargo in vacuolar macropinosomes which are also hubs for receptor signalling and for endocytic and recycling membrane traffic (Stow, Hung, & Wall, 2020; Swanson, 2008).

Recent developments in microscopy include the introduction of the lattice light sheet microscope (LLSM) (Chen et al., 2014) which utilises a 2D optical lattice of Bessel beams to achieve resolution in X, Y and Z plane near diffraction-limited level with high signal-to-noise ratio. Beside maintaining good optical resolution, another key advantage of the LLSM is the low phototoxicity and photobleaching of specimens, which enables cells to be surveyed for extended periods of time in physiologically relevant conditions. This imaging now enables live, 3D fluorescence imaging of sufficient speed, duration and temporal-spatial resolution to adequately capture and record exquisite details of dynamic cell surface projections. LLSM studies have recorded novel features of ruffling, macropinocytic cups, filopodia and other surface projections in amoeba and vertebrate immune cells (Condon et al., 2018; Fritz-Laylin et al., 2017; Manley et al., 2020; Veltman et al., 2016). LLSM recordings in 3D extending over many hours can capture thousands of cell surface protrusions., routinely resulting in terabyte-scale datasets that cannot be interrogated manually or with traditional segmentation, nor with techniques such as thresholding and automatic spot detection, which require careful calibration or manual editing (Heddleston & Chew, 2016). Machine learning provides a promising approach, allowing manually defined example data to be extrapolated via sophisticated models. These models can be applied at scale, robustly performing tasks such as classification and semantic segmentation.

We earlier used live imaging and LLSM to record cell surface ruffling on activated macrophages (Condon et al., 2018). The macrophage surface has many spike-like projections or filopodia, in addition to constant, undulating wave-like membrane ruffles (Swanson, 2008). LLSM also revealed a new type of ruffle, so-called ‘tent pole ruffles’ characterised by filopodia (tent poles) embedded in the ruffles. The tent poles appear to raise up the intervening ruffle and then twist together to close the ruffle for the formation of fluid-filled macropinosomes. The distinctive tent pole ruffles were characterised on lipopolysaccharide (LPS) activated macrophages but are also detected on other cell types such as cancer cells (Condon et al., 2018). The relationships between filopodia, ruffles and tent pole ruffles remain to be fully investigated to define their modes of formation and deployment by cells under different conditions. These and other cell projections perform key roles in cell migration, immune defence, and the formation of macropinosomes for environmental sampling and nutrient acquisition. LLSM datasets are valuable troves of data for the quantitative analyses required to gain critical insights into cell surface behaviours, but machine learning is necessary to unlock this information.

Here we set out to develop machine learning software for the analysis of actin-rich protrusions consisting of spike-like filopodia, ruffles and tent pole ruffles on the surface of Lifeact-labelled, activated macrophages in LLSM datasets (Fig.1A). We demonstrate a scalable, configurable and modular analysis pipeline suitable for large 4D microscopy datasets (Fig. 1B), LLAMA (**l**arge scale **l**ight microscopy **a**nalysis with **ma**chine learning). All code is provided (github.com/jameslefevre/4D-microscopy-pipeline, github.com/jameslefevre/visualiser-4D-microscopy-analysis), as well as a detailed set of protocols (see supplementary material). The system is designed to identify and track cell surface projections in such datasets, but it may be readily adapted to other types of large 4D image datasets. The pipeline is based on a machine learning approach to perform semantic segmentation, assigning each voxel to a defined class, followed by object separation and tracking algorithms designed to deal effectively with ambiguous structure delineation. The output datasets contain rich information on cell surface features over time, suitable for both statistical analysis and visualisation. This pipeline also provides a template for the application of ImageJ (Schneider, Rasband, & Eliceiri, 2012) based algorithms to large scale image datasets, including deployment to HPC (high performance computing) systems. We also provide a 4D visualisation tool, the LLAMA visualiser, designed to support the pipeline (training data selection, parameter selection and validation) as well as to visualise the outputs.

**Figure 1.**
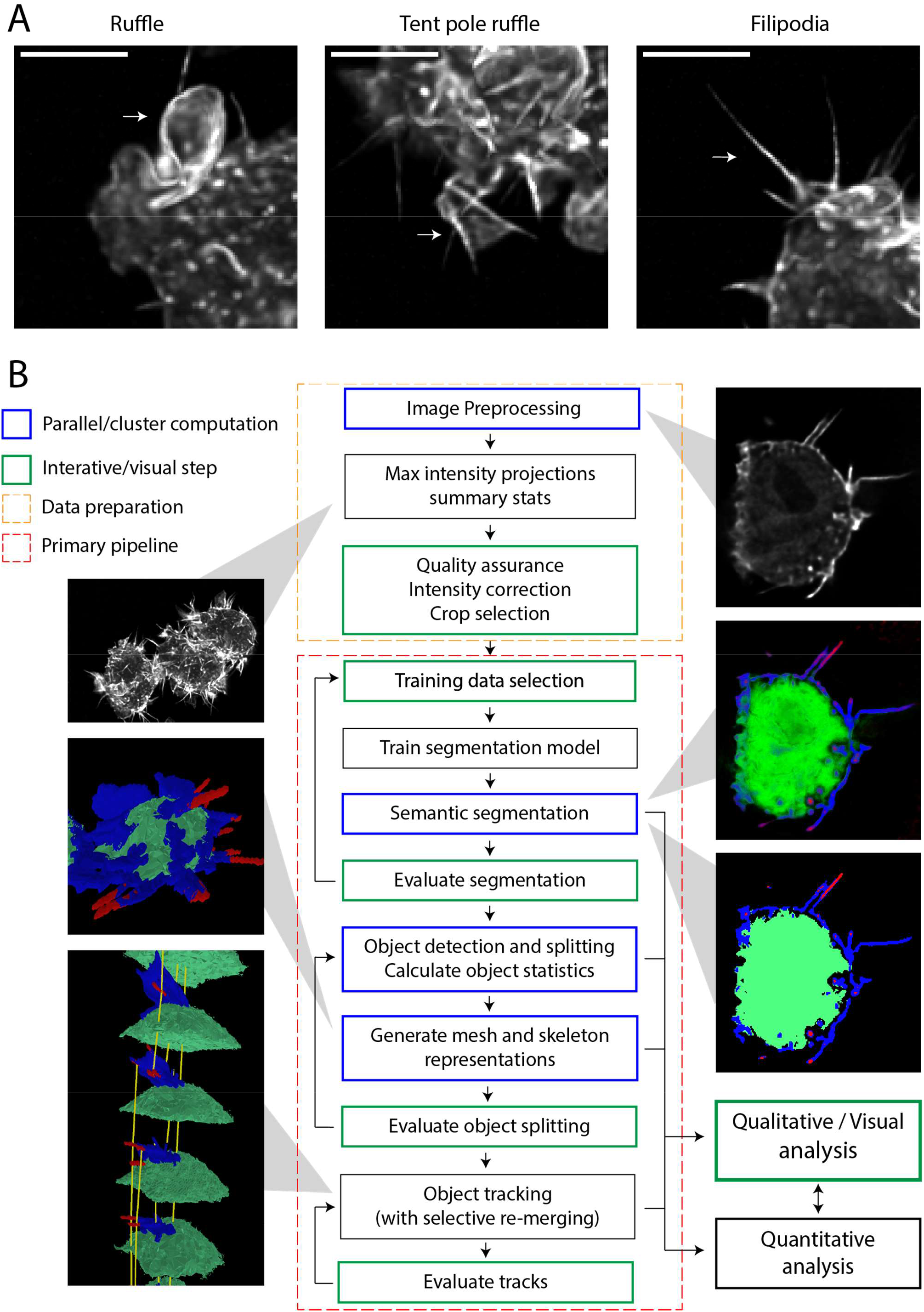
**(A) Macrophage membrane projections** Ruffle, tent pole ruffle and filipodia. Example images from LPS treated cell samples, maximum intensity z-projection. Scale bars are 5μm. **(B) Overview of the LLAMA image analysis system** Key steps in analysis pipeline; images are illustrative. Our system is designed for systematic analysis of large scale 4D microscopy datasets and provides high throughput by performing intensive computations (blue) in parallel for each time step, suitable for an HPC cluster. The LLAMA visualiser is designed to support all interactive steps (green), including quality assurance, segmentation model development and parameter selection as well as analysis of results. Development of the segmentation model and the selection of parameters for object detection and tracking should be performed using a representative selection from the dataset. The pipeline can then be run non-interactively, using uniform settings to ensure comparable output across the dataset. See Methods and Materials and supplementary protocols for details.

Segmentation classes used to dissect this imaging were background, cell body, ruffle and filipodia, where filipodia include the recently discovered “tent pole” structures contained within some ruffles (Condon et al., 2018). This was used to quantify the effect on macrophages subjected to two different interventions that stimulate ruffling: i) bacterial lipopolysaccharide (LPS) LPS is a potent endotoxin that stimulates morphological changes associated with arming innate immune responses in macrophages through actin reorganisation and tyrosine phosphorylation of Pyk2 and focal adhesion, paxillin (Williams & Ridley, 2000); and ii) macrophage colony stimulating factor (CSF-1), a cytokine involved in differentiation of macrophages that induces cells via WAVE2-Abi mediated pseudopod assembly for cell chemotaxis (Kheir, Gevrey, Yamaguchi, Isaac, & Cox, 2005). Both stimuli also induce macropinocytosis, an actin driven process that facilitates the bulk engulfment of extracellular fluid via ruffling (Canton et al., 2016; Zanoni et al., 2011). For these studies, samples containing 17 complete cells were each imaged at a resolution of 1.04μm × 1.04μm × 2.68μm × 5.3s for 53 minutes total over two captures (before and after treatment). Cell tracks were matched between the two captures and excluded if cells moved partially or fully out of the field of view during imaging. 901,696 objects were found that were associated with the analysed cells, arranged into 76,386 tracks that met minimum size thresholds. Statistics produced included volume, maximum extension from the cell surface and the set of adjacent structures. We identified and analysed 1,188 significant ruffling events, defined by peak ruffle volume exceeding 14.5μm^3^.

## Results

### Semantic segmentation

Our semantic segmentation approach is based on the Trainable Weka 3D software (Arganda-Carreras et al., 2017), which is implemented as a plugin to the ImageJ image processing platform (Schneider et al., 2012). This machine learning tool produces segmentations using a two-step process. Using ImageJ, a range of sophisticated pre-defined 3D image features are computed at multiple scales, capturing rich spatial context for each pixel. Then the set of features for each pixel is used with a selected machine learning algorithm from the Weka toolkit (Witten, Frank, Hall, & Pal, 2016) to produce a per-pixel classification. We employed a reengineered high-throughput pipeline suitable for use on a large scale 4D dataset with cluster-based computation (github.com/jameslefevre/4D-microscopy-pipeline). See Methods and Materials for details.

Training data was obtained from four image stacks, selected from four different captures and two imaging sessions. Compared to the deep learning models sometimes used for semantic segmentation (Zinchuk & Grossenbacher-Zinchuk, 2020), the Trainable Weka approach requires only a very small selection of training data, although high quality results may require an iterative process involving assessment of representative segmentations, followed by updating the model with additional training data. A total of 16330 voxels were labelled with one of the 4 segmentation classes, equivalent to less than 0.1% of a single image stack cropped to the size of a macrophage (approx. 60000μm^3^). Two revision steps were used to refine the model and adjust for heterogeneity in noise and contrast between image captures. Fluorophore intensity was adjusted between and within captures using cytoplasm intensity as a benchmark and modelling an exponential decay curve in each capture; the same intensity adjustment process was also used for analysed data. In addition, the segmentation model was made more robust against base fluorophore intensity variation by 4-fold replication of training data with intensity scaled by factors 0.5, 0.75, 1, 1.25. The final model used the Weka random forest algorithm with default parameters on class-balanced training data. The random forest was selected from the range of models available in Weka as it was judged to perform best in generalising beyond the training data. The full range of features was used, with kernel size (sigma) of 1,2,4,8, and 16μm (16μm is approximately 15×15×6 voxels). For computational efficiency, a 2-fold down-sampling in x and y was used for all features at sigma=16, and the ImageJ filters (mean, median, minimum, maximum, variance) at sigma=8. The maximum sigma value must reflect the amount of spatial context necessary to classify each voxel, and the down-sampling feature was added to our system to minimise the computational cost of computing features at larger scales. The wide range of features used was necessary to provide robust and clean segmentations in the presence of heterogeneous image quality and noise; otherwise a more parsimonious model could have been used, allowing faster computation.

Model validation on the broader image data necessarily relies on user judgement, reinforcing the need for effective visualisation. We considered producing test data in the form of full stack manual segmentations, but truly objective and unambiguous segmentation was found to be impossible on our data at voxel resolution: producing clean boundaries using a consistent model is more useful than replicating an imperfect manual boundary.

The robustness of the model was tested on the training data using a 10-fold cross validation, giving a 0.73% error rate. Training on similar image datasets consistently gave cross validated errors below one percent, and this continuing very low error rate shows a highly robust model.

The segmentation model was successful in identifying the four classes, distinguishing ruffles and filipodia and identifying filopodial-like “tent poles” embedded in major ruffle structures (Fig. 2).

**Figure 2:**
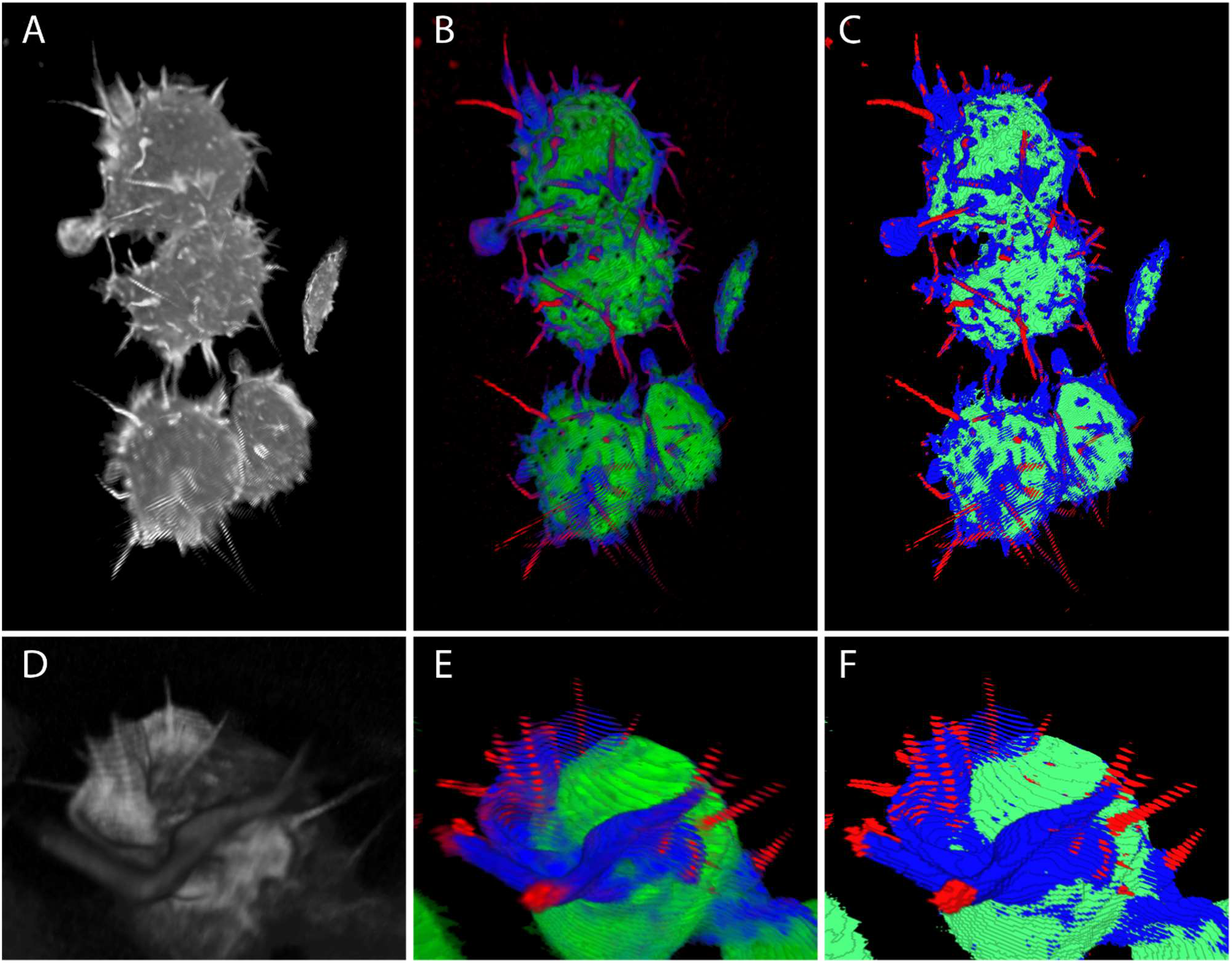
Segmentation. Sample segmentation output, showing macrophage cells imaged on the LLSM, segmented into background (black/transparent), cell body (green), ruffle (blue), and filipodia/tent pole (red). (ABC) Full image stack, view rotated so that plate is on left side. (DEF) Detail of ruffle containing tent pole like structures. (AD) Original image after pre-processing (de-skew and deconvolution). (BE) Probability map, showing the estimated probability that a voxel will occur in each class. (CF) Semantic segmentation; each voxel is assigned to the single class considered most likely. All images produced using our LLAMA visualisation software (github.com/jameslefevre/4D-microscopy-pipeline).

The segmentation results, displayed alongside the LLSM imaging using LLAMA visualiser, proved effective in helping to manually identify tent pole ruffling events (Videos 1,2).

#### Video 1,2: Tent pole ruffling

Two selected examples of tent pole ruffling events in LPS treated macrophage cells, showing LLS imaging and corresponding segmentation probability map. Segmentation colours are as described in Fig. 2. Videos show direct capture from the LLAMA visualiser with no additional image processing.

### Object detection and tracking

Semantic segmentation is designed to distinguish between classes of structures, and it will not necessarily separate individual objects. However, the most useful quantitative analysis of cell imaging will typically be based on individual cells or subcellular structures, tracked over time. In Fig. 2C we can clearly distinguish 4 cells (and the edge of a structure which is primarily outside the imaging frame), but these are not fully separated in the segmentation. These cells must be delineated, and each tracked over time. Macrophage ruffles are more complex and highly variable, representing perhaps the worst-case scenario for object delineation and tracking. It can occasionally be unclear even to the human analyst whether an object should be considered as one structure or two, or where the boundary is, or when an object should first be considered a ruffle that is distinct from the cell membrane. Biological variation and noise in the imaging process mean that an automated process attempting to decide these questions without putting the data in temporal context may give inconsistent results over a sequence of time steps, making coherent tracking of structures impossible. However, due to the scale of the data and the need for computational feasibility it is necessary to *segment* each image stack in isolation, without incorporating information from adjacent time steps, and the algorithm used for this task cannot rely on manual editing.

In response to these challenges, we developed a sophisticated and configurable approach in which touching structures are split using a watershed algorithm (Beucher & Lantuéj, 1979) on edge distance (see Methods and Materials for details), then selectively recombined as part of the tracking process in order to pool information across time steps and produce coherent tracks of structures over time (Fig. 3). The computationally expensive watershed split step is performed in parallel for each image stack prior to the tracking algorithm, consistent with a scalable high-throughput analysis pipeline. Methods involving a re-merging step in combination with watershed split are well known, for example (Wählby, Sintorn, Erlandsson, Borgefors, & Bengtsson, 2004). The key innovation in our algorithm is to integrate this re-merging with the tracking algorithm as a way of efficiently pooling information across time. The algorithm is described in detail in Methods and Materials.

**Figure 3:**
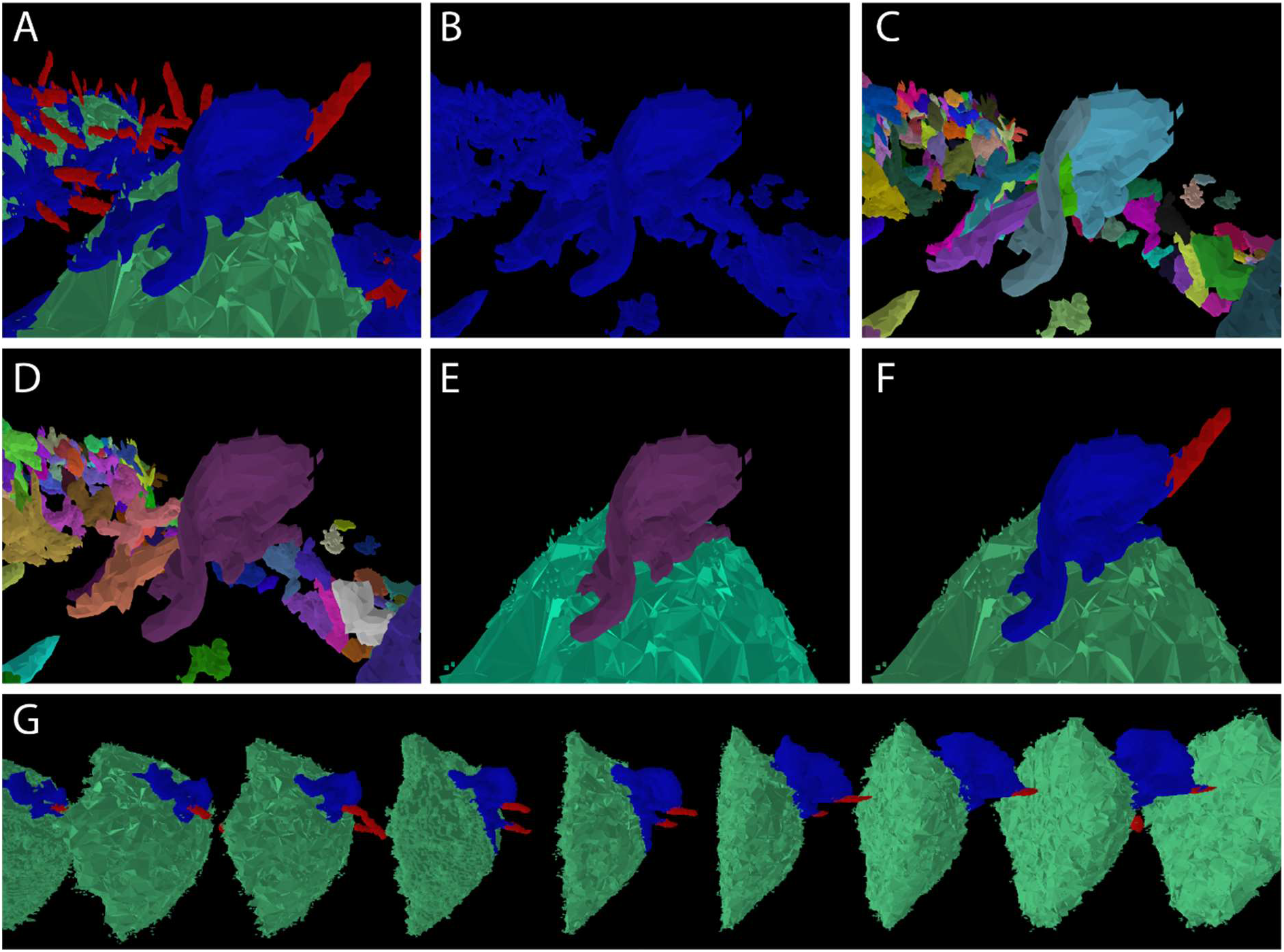
Structure delineation and tracking. (A) Semantic segmentation into cell body (green), ruffle (blue), filipodia/tent pole (red), with a prominent ruffle in the foreground. (B) Isolation of ruffle class. (C) Watershed split algorithm separates ruffle from touching objects but incorrectly splits the foreground structure; except in the simplest cases, this initially excessive splitting is necessary to ensure that all required separations are performed. (D) Structure boundaries after re-merging step; this is integrated with the tracking algorithm to maximise consistency with other time steps. (E) Foreground ruffle correctly delineated and associated with the cell body. (F) Ruffle with associated call and filipodia structures (identified using adjacency data) using the original class colour scheme. (G) Structures in F tracked over time; the timestep shown in A-F is in the central position. All images shown were produced using our LLAMA visualiser using only the standard display options available in the graphical user interface, with no post-editing. This application provides a powerful tool for customisation and validation of the algorithm as well as visualising results.

A potential challenge with the macrophage segmentation is a degree of unresolvable ambiguity between the ruffle and filipodia classes; some structures appear to be truly intermediate, giving complex and unstable segmentations. In order to allow analysis of these structures, such as filipodia in the process of forming or decaying, we also applied the object detection and tracking algorithm to the merged ruffle and filipodia classes, in addition to the separate analyses. The average class probabilities of each structure are provided as an additional metric (measuring the degree to which a composite structure most resembles a ruffle or filipodia). This capacity to analyse merged segmentation classes is included as a general feature in the protocols and code provided.

### Statistical analysis

Our system provides a rich set of data for each structure, including size, position, shape, orientation and distance from a specified reference class (measuring, for example, the maximum extension of a structure from the cell surface). The contact between touching objects of all classes is quantified, allowing relationships between structures to be determined. These data are recorded in tabular form, for easy loading into any statistical software. In addition, skeleton and mesh representations of objects are (optionally) produced. These are intended as an aid for visualisation using the LLAMA visualiser or other software, and may also be used to provide additional statistical information such as the length of linear structures.

Each macrophage sample analysed was imaged for 300 timesteps before and after treatment, at intervals of approximately 5.3 seconds; we refer to these as the pre and post captures. Matching of cell tracks allowed direct pre vs post comparisons for each cell after stimulation. Observed changes were compared to untreated control cells, which were prepared in parallel and imaged in the same session. Analysis was performed in the R statistical programming environment using the track files produced at the final stage of the image analysis.

Many ruffles and filipodia were found to extend along the slide surface and appeared to be attached to the slide, resulting in a morphology distinct subset from the structures in the upper region of the cell. Differential behaviour was also considered likely between these regions, so for analysis we classified structures as slide-proximal or slide-distal, depending on whether the distance from the estimated slide position to the structure centre of mass is less or greater than 1.5μm.

In the following sections we demonstrate types of analysis and conclusions that can be drawn using these methodologies. Distinct patterns of macrophage ruffling from LPS and CSF stimulation (as discussed in the introduction) were seen, with increased activity in the distal and proximal regions of the cell surface respectively. Using data at the level of individual tracked structures, we could attribute this to increased frequency of ruffling events with CSF, while LPS simulation lead to larger as well as more numerous ruffles. The duration of individual ruffling events did not change significantly in either case.

### Distinct patterns of increased ruffling and filipodia volume in LPS and CSF treated cells

We performed an analysis based on the total volume of ruffle and filipodia structures in each cell before and after cell treatments (Fig. 4), identifying spatially distinct patterns of increased activity. A consistent increase in ruffle volume was seen for both LPS and CSF cells, however this increase occurred exclusively in the plate-distal regions of the LPS treated cells, and the plate-proximal regions of the CSF treated cells. In contrast, a consistent increase in filipodia/tent pole volume was seen in both proximal and distal regions of the LPS cells, while no significant changes were seen in the CSF treated cells.

**Figure 4.**
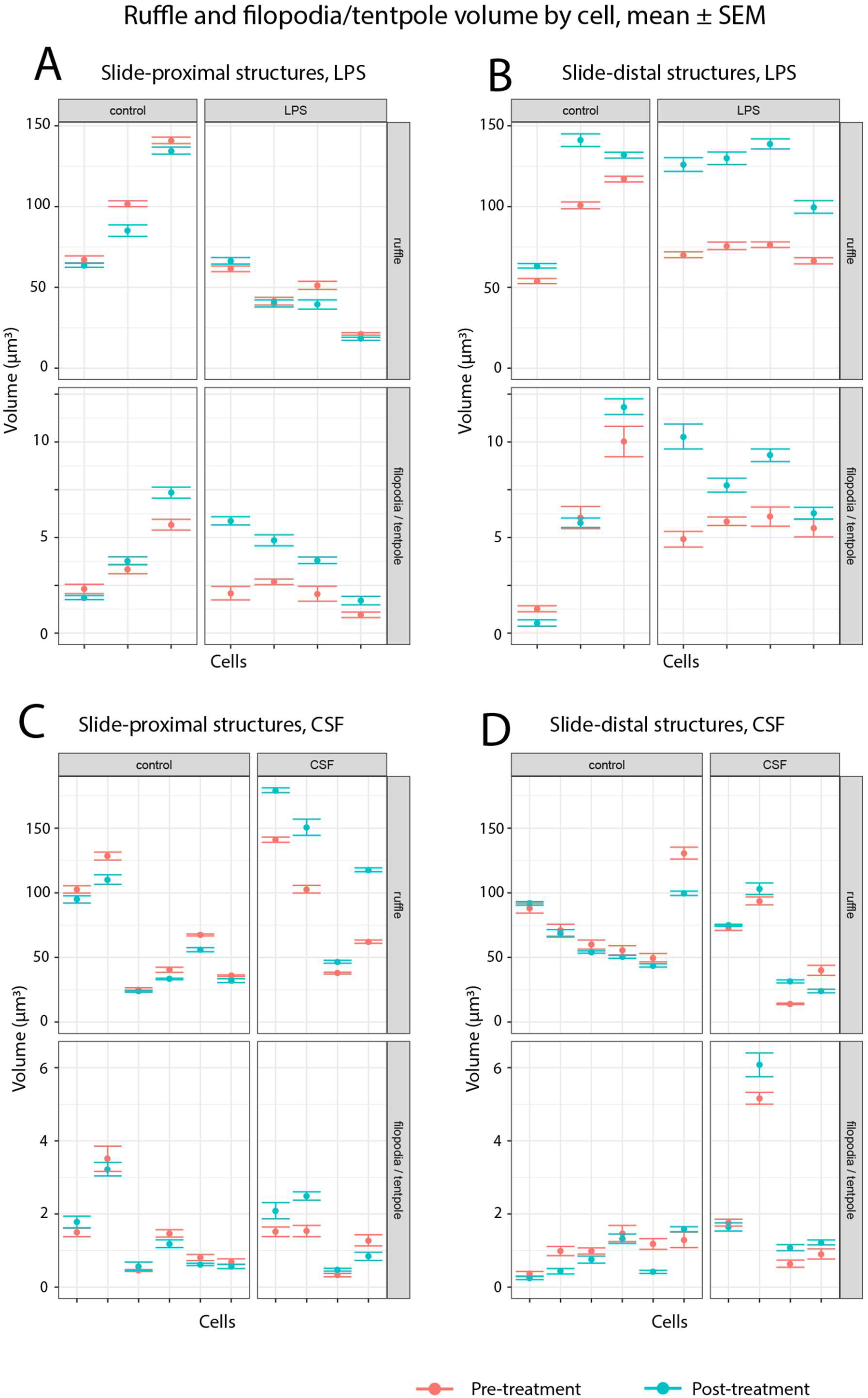
Ruffle and filipodia volume. Aggregate ruffle and filipodia (including tent pole) volume per cell, mean and SEM pre and post treatment. Structures are classified as plate-proximal (within 1.5μm of imputed plate position) or plate-distal (greater than 1.5μm from plate).

(A) LPS experiment, plate-proximal; (B) LPS experiment, plate-distal; (C) CSF experiment, plate-proximal; (D) CSF experiment, plate-distal. Means are calculated across the time steps in each capture, with SEM adjusted for autocorrelation. This adjustment does not account for possible trends over time that are independent of treatment, so changes in control cells (left) are included for comparison with treated cells (right). In the LPS treated cells, consistent increases are seen in the slide-distal ruffles and the filipodia in both distal and proximal regions. In contrast, the CSF experiment exhibits consistent increase in the slide-proximal ruffles and no change to filipodia volume in either region.

### Increased ruffling associated with higher frequency of events in CSF cells, increased frequency and size in LPS cells

The aggregate volume analysis in Fig. 4 demonstrates spatially biased patterns of increased ruffling. An important question is whether this is reflected in any change to the nature of the ruffling events that are associated with macropinocytosis or cell motility under different conditions of cell stimulation. As from our data and others (Barthwal et al., 2013; Canton et al., 2016; Lin, Mintern, & Gleeson, 2020; Norbury, Hewlett, Prescott, Shastri, & Watts, 1995; Pixley, 2012), CSF regulates mesenchymal cell motility and macropinocytosis, whereas LPS facilitates primarily macropinocytosis in macrophages under these conditions. Therefore, is the increased aggregate volume associated with an increased number, size or duration of events during these different processes? The track dataset provides a rich set of information for addressing questions such as these. Individual ruffle tracks could not be reliably used as a proxy for ruffling events, as numerous tracks were identified which persisted well beyond a distinct ruffling event and even included two or more consecutive events; this is consistent with actin recycling between ruffling events, and with the spatial clustering behaviour observed in (Condon et al., 2018). We isolated major ruffling events from the tracked ruffle structures by identifying track intervals in which the peak volume is above 14.5 μm^3^ (5000 voxels; this threshold was established by inspection in the visualiser) and the volume at the end points is less than half the peak value. Hence, we are able to classify each LifeAct-labelled event from the initiation of actin polymerisation/extension (increased in ruffling volume) until its retraction.

More ruffling events were observed after treatment in both LPS and CSF cells, with the increases occurring in the slide-distal region of the LPS cells and the slide-proximal region of the CSF cells (Fig. 5A), consistent with the aggregate volume results. There was also an increase in the median peak volume of the slide-distal ruffling events in the LPS cells (Fig. 5B). We observed that LPS stimulate circular dorsal ruffles that protrude exclusively from the plate-distal region/or the peripheral surface of the cell. Whereas in CSF stimulated cells, lamellipodia-like structures along the plate proximal region predominantly emerge upon stimulation. Filopodia/tent-poles, as finger-like linear actin structures responsible for sensing chemical and mechanical cues, elongate and increase in quantity under both circumstances.

**Figure 5:**
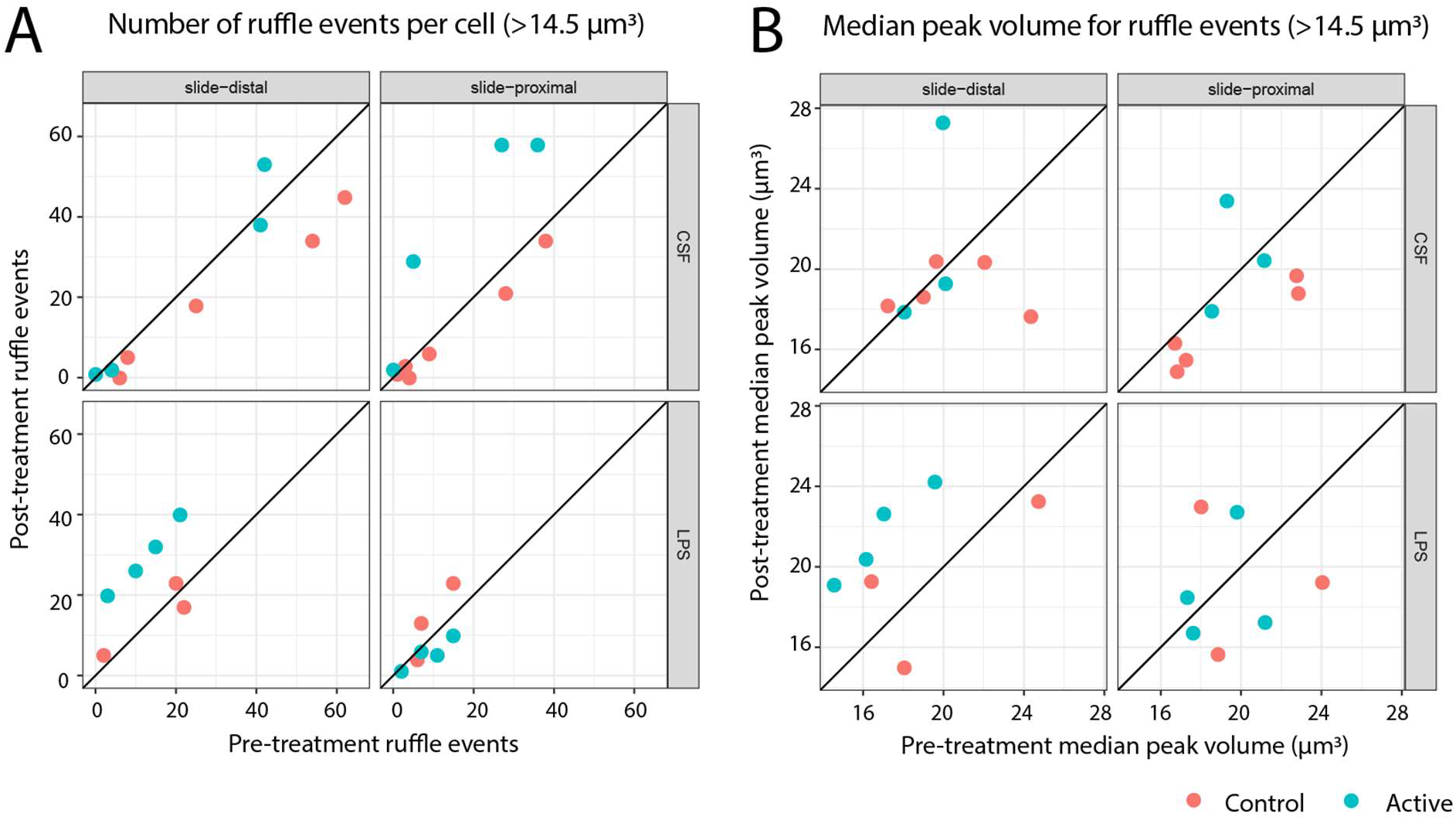
Ruffling events. The number of ruffling events increases in the slide-distal region of LPS treated cells and the slide-proximal region of CSF treated cells, while peak volume increases for slide-distal LPS only. Ruffling events are defined as a ruffle structure tracked over a time period in which the peak volume is at least 14.5 μm^3^ (5000 voxels) and the volume at the start and end of the time period is less than half this peak value (or the end of the capture period is reached). (A) Number of ruffling events per cell, before and after treatment. (B) Median peak ruffle size per cell, before and after treatment. Points above the diagonal line indicate an increase in number of events or median size.

A range of additional metrics are automatically generated by our system, and several were analysed but not included here, as no clear experimental effect was found. Duration of ruffling events was considered as a possible contributing factor to the overall increase in activity, but no change was detected for either CSF or LPS, suggesting that ruffle deployment but not structure is modulated in different physiological conditions. The maximum extension of each structure from the cell surface was measured, and the plate-proximal structures were found to be substantially more elongated on average, but this difference did not depend on treatment. Further null results were given by measurements of filipodia size and the number of filipodia associated with each ruffle.

### Tent pole ruffling motifs occur within a diversity of ruffling behaviour

We are interested in whether tent pole ruffling is a common event. The plate-distal regions in the LPS treated macrophages showed a high level of ruffle and filipodia/tent pole activity (Fig. 4), and examples of filopodia ruffling were readily found (Videos 1,2). However, an initial visual assessment did not find that the majority of ruffling events (Fig. 5) corresponded directly with tent pole ruffling, although apparent tent poles were often present, and aspects of the previously described behaviour could be identified. This visual assessment was greatly complicated by the rapid and dynamic nature of actin turnover.

To gain a more systematic understanding we undertook a complete analysis of the first 13 minutes (150 timesteps) in an LPS treated sample containing 4 macrophage cells.

Firstly, we manually identified four stereotypical examples in which (at a specific time) a pair of prominent tent poles were joined by a ruffle “veil”, and analysed these examples using the object statistics to develop a filter to detect all similar cases. The selected filter required: (1) a ruffle object of volume 10.15μm^3^ (3500 voxels) or greater, with at least 2μm mean and 4μm maximum distance from the cell surface, and at least 1.5μm mean distance from the plate; (2) two filipodia/tent pole objects adjacent to the ruffle, each with volume 0.22μm^3^ (75 voxels) or greater, and minimum contact score with the ruffle of 240 (this arbitrary score is based on pairs of adjacent voxels, one in each object).

Computationally applying these filter criteria to the 4 macrophage cells over 150 timesteps, we identified 133 examples belonging to 33 distinct tracked ruffles. We then sampled 10 of these tracked ruffles at random for detailed visual analysis. In 9 cases we observed rapid twisting of the tent pole pair during or after the collapse of the connecting ruffles, which we consider the canonical feature of tent pole ruffling and a key closure mechanism for macropinsomes (Condon et al., 2018). Variations in the origin and timing of the ruffles and tent poles were also recorded as follows. Ruffles developed from a) part of a pre-existing ruffle structure (4 cases, see Video 4); b) de novo on the dorsal region of the cell (3 cases, of which only 1 features tentpoles during the initial emergence, see Video 3); c) de novo on the proximal region of the cell, attached to the plate(2 cases). Tent poles appeared either during the terminal phase, after the ruffle formed a semi-circle and started to close (5 cases, see Video 3); tent poles exhibited a configuration with a “veil” in between forming prior to the final phase (3 cases, see Video 4) and a tent pole pair appeared to emerge from the hinge point of a large ruffle (1 case, Video 5).

Our approach of computational filtering combined with visual analysis allows us to detect different features within a range of complex and multi-facetted ruffles, including distinctive tent pole ruffling events. Importantly, the variety of behaviours observed here suggests that tent pole ruffling exists on a continuum with non-tent pole ruffling. The fact that tent pole ruffling and the involvement of filipodia in ruffle formation are increased by LPS activation of macrophages (Condon et al., 2018), suggests that this continuum of ruffling morphologies is ‘tunable’. By revealing that ruffling constitutes a range of membrane formations, rather than discrete subtypes of ruffles, now frames future studies that will dissect the physiological demands and molecular functions that drive this variation.

#### Videos 3-5

Representative examples selected from ten randomly sample ruffling events that include a “tent pole ruffle” configuration in which a pair of filipodia / tent poles are connected by a prominent ruffle. Segmentation colours are as described in Fig. 2. Videos show direct capture from the LLAMA visualiser with no additional image processing.

## Discussion

Realising the full potential of terabyte scale microscopy data requires new approaches to image analysis. Even storing and transferring image datasets on this scale can overwhelm the local IT resources available to a typical researcher; a high-end workstation running specialised software is certainly capable of visualising an individual image stack produced by an LLS microscope, as well as performing analytical processes such as thresholding and object counting, but running a systematic quantitative analysis over hundreds of time steps is not feasible. Automated processes running on high performance computing facilities are required, and the algorithms employed must be robust enough to produce reliable results with minimal recourse to manual editing and adjustment. Machine learning methods provide a promising approach and have proved capable of producing robust semantic segmentations of microscopy imaging. Segmentation into defined tissue classes requires supervised machine learning, meaning that data must be labelled for training, and the models derived from this training data represent a sophisticated means to extrapolate to the larger dataset. But such a model can only be reliably applied on data that is well represented by the training data selection, and may be invalidated by changes in experimental conditions, markers and microscopy settings. In response to these challenges, we selected a machine learning approach featuring rapid model training with minimal data labelling requirements. We especially emphasised the development of a custom visualisation tool, to optimise training data selection and the assessment of draft segmentations.

The LLAMA image analysis system was designed to automatically detect and decipher characteristics of cell membrane protrusions from large scale data acquired through the LLSM, with integration of statistical and visual analysis. Semantic segmentation provides the starting point for an object delineation and tracking system designed to convert the segmentation into a rich set of data at the level of individual structures over time. While designed as a general-purpose tool, our primary goal was to analyse macrophage ruffles, filipodia and the filipodia-like “tent poles” embedded within ruffles. The complex behaviour of macrophage ruffles proved a particular challenge, and the system features a novel approach designed for separating and tracking these structures, under the constraint that segmentation and other computationally difficult tasks are performed for each time step in isolation. The algorithm is parameterised to allow customisation to other structure types and was successfully adapted to the cell body and filopodia classes. Our system is designed for flexible deployment and is suitable for cluster or cloud computing, providing a general-purpose system for tracking and quantifying structures in large scale 4D microscopy data. The included code also provides a template for deploying an ImageJ based processing pipeline on high performance computing facilities.

Using this system, we were able to perform for the first time a systematic quantitative analysis of RAW264.7 macrophage cells under two experimental conditions, LPS and CSF treatment. We were able to tease out features that varied under different stimulation; these included the frequency of occurrence, location (proximal or distal to plate) as well as size of the actin structures on the cells. In addition, there are aspects of these actin structure where no differences were found, such as the mean lifetime and maximum distance from the cell surface. These physical variations in actin structures reflect the different physiological requirement of the cell under LPS (pathogen uptake) versus CSF (cell migration and invasion).

We were also able to conduct a systematic qualitative analysis of tent pole ruffling in LPS treated cells, using the capacity of the LLAMA visualiser to link numerical and visual data. This analysis highlighted the considerable challenge of completely characterising macrophage membrane protrusions, with the largest ruffles in particular exhibiting complex and multi-facetted behaviour. Most importantly, we were able to demonstrate continuity between tent pole ruffling and the wave-like ruffling behaviour as traditionally understood.

## Methods and Materials

### Cell culture

The RAW 264.7 macrophage-like cell line was obtained from ATCC, maintained regularly by passage in RPMI 1640 medium (Sigma Aldrich) supplemented with 10% heat-inactivated FCS (Interpath services) and 2mM L-glutamine (Invitrogen) at 37°C, 5% CO2. Cells were routinely checked for mycoplasma by PCR. RAW cell lines stably expressing LifeAct were generated using a PEF6_GFP_LifeAct plasmid and maintained under Blasticidin (ThermoFisher) selection.

### Cell imaging on the Lattice light sheet microscope

RAW 264.7 cells stably expressing GFP-LifeAct were plated at 0.15 × 10^6^ cells/ml on 5mm coverslips one day prior to the experiment. For LLSM imaging, cells were matched-paired and incubated in CO^2^-free conditions in L-15 medium (supplied with 10% FCS 2mM L-glutamine and 10μg/ml Pen/Strep) and LPS (Sigma-Aldrich) or CSF-1 (Miltenyi Biotec Australia Pte Ltd), were added at 100ng/ml. Cells were imaged for 20 mins with and without these activators at 20ms exposures, for at least 121 planes. The imaging was carried out using a 3i lattice light sheet microscope with excitation by Coherent Sapphire 488nm and MPB Communications 560nm diode lasers at 1-2% AOTF transmittance through an excitation objective (Special Optics 28.6 × 0.7 NA 3.74-mm water dipping lens) at an angle of 31.8 degrees. Individual sheets were generated using 3i Slidebook software (488nm 52 beams, 0.960mm spacing, 560nm 45 beams, 1.102 mm spacing both with a cropping factor of 0.150mm and spacing factor of 0.970) through a 0.550/0.493 NA annular mask. Fluorescence signal was detected with a Nikon 25x 1.1 NA CFI Apo LWD objective and 2.5X tube lens (62.5x total system magnification) using 2x Hamamatsu Orca Flash 4.0 V2 sCMOS cameras running SlideBook 6 software (3i, Colorado, USA).

### Image pre-processing

LLSM image captures were uploaded to a file-storage server as a sequence of image stacks in tiff format (one image stack volume per time step). These were de-skewed to correct for the slanted image capture used, then deconvolved using the Microvolution software (50 deconvolution steps). An initial review was performed using maximum intensity projections in the y and z dimensions, assessing image quality and completeness, and determining where image stacks could be safely cropped. For each usable image capture, an intensity adjustment factor was calculated by comparing the average initial cytoplasm intensity to a reference value (600). The cytoplasm intensity was estimated by applying a gamma=0.5 transform to the set of voxel intensities to moderate the dynamic range, then finding the second histogram peak, confirmed using the area measuring tool in ImageJ. In addition, an exponential decay curve was fitted to the mean intensities of the image stacks in each capture (plotted and reviewed for goodness of fit). This provided a per-time step intensity adjustment which was used in combination with the initial intensity adjustment prior to semantic segmentation. Note that these pre-processing steps are not part of the LLAMA image analysis system that we are presenting and are not included in the protocols, although intensity adjustment and cropping parameters may be used with segmentation, and for convenience the code repository includes the script used to produce maximum intensity projections and image stack summary stats (max_proj_and_summary.groovy).

### System overview

Our computational pipeline is provided as a set of linked protocols (see Supplementary material), with complete and commented code (github.com/jameslefevre/4D-microscopy-pipeline). The implementation primarily uses headless ImageJ (Schneider et al., 2012) processes, invoked via parameterised Groovy scripts, which combine core ImageJ functionality with selected plugins and custom extension code. This provides a common interface to each computational step with flexible calling options, including interactively from the ImageJ interface, as a background process on a local machine, or on a remote server or cloud service. While this approach exposes some complexity, requiring manual editing of scripts and selecting paths and other parameters, it also provides extensive flexibility in deployment, including local or remote execution as well as modification, removal or substitution of steps. The protocols provide a detailed guide for deploying the pipeline using a remote cluster with a PBS batch job system to perform the main computations (semantic segmentation and object analysis, steps 2-5 in table below) in parallel for each time step, enabling a scalable, high-throughput system. The code provided includes example PBS job scripts. ImageJ/FIJI was selected since it is free and open source, provides access to a wide range of image processing algorithms (via plugins as well as core ImageJ), and also has the benefit of a fully cross-platform Java-based system and simple installation (admin privileges not required, optionally bundles Java to avoid system dependency).

The custom 3/4D LLAMA visualisation software we developed (github.com/jameslefevre/visualiser-4D-microscopy-analysis), is built on the Processing 3 environment and language (Reas & Fry, 2006), which provides a powerful framework for interactive visualisation; this is the only component of the system which is not based on ImageJ. It should be noted that the visualiser is not directly part of the analysis pipeline, and following our modular approach it may be ignored, or alternative tools used. However, we found this to be a vital support tool throughout the process, enabling easy and direct comparison of imaging with segmentations and object representations, and between alternative versions of these outputs. This tool is designed specifically to support training data selection in 3D and customisation and quality control across the pipeline, as well as visualising outputs. It features rapid switching between slice and 3D view, and between the original image and one or more, from a single selected perspective. The segmentations may similarly be compared to the object representations derived from them. The protocols document includes a guide to using the visualiser as part of our system, and a manual is also included with the code. The supplementary material includes videos demonstrating its use and key features (visualiser_demo1_training_data_selection and visualiser_demo2_object_and_track).

**Table 1:**
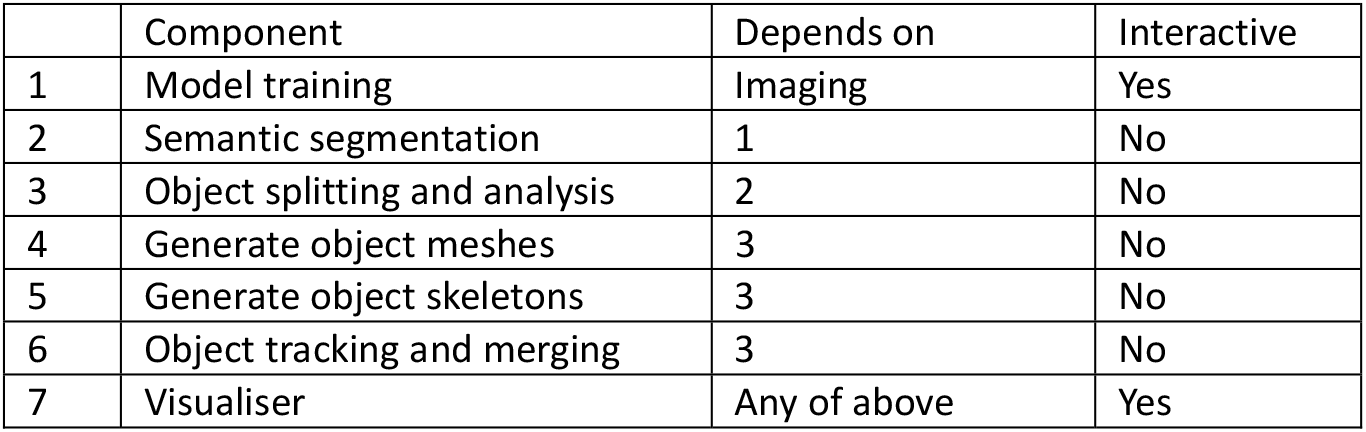
LLAMA System components.

|  | Component | Depends on | Interactive |
| --- | --- | --- | --- |
| 1 | Model training | Imaging | Yes |
| 2 | Semantic segmentation | 1 | No |
| 3 | Object splitting and analysis | 2 | No |
| 4 | Generate object meshes | 3 | No |
| 5 | Generate object skeletons | 3 | No |
| 6 | Object tracking and merging | 3 | No |
| 7 | Visualiser | Any of above | Yes |

**Table 2:** Tracking parameters.

| Parameter | Name in script | Purpose |
| --- | --- | --- |
| $W_l$ | logSizeWeight | Weight to put on size difference when matching objects |
| $d_{max}$ | matchThreshold | Maximum distance for matching |
| $Q_p$ | relativeNodeDistance_referenceValue | Calibrates the object merging metric based on centre of mass proximity |
| $Q_c$ | relativeNodeContact_referenceValue | Calibrates the object merging metric based on contact between objects |
| $W_c$ | relativeNodeContact_weight | Relative weight for contact metric versus distance metric |
| $W_{match}$ | matchScoreWeighting | Relative weight on matching metrics versus merging metrics |

### Semantic segmentation

The protocols document provides detailed instructions for the segmentation model training process and for the deployment of trained models. It is often necessary to modify the model one or more times after evaluation of segmentation results on a larger set of image data, and an additional protocol is provided as a guide for this iterative process.

Our machine learning segmentation approach is based on the Trainable Weka software, and we briefly describe this system and our reengineered pipeline, designed to more effectively deal with large scale data. The Trainable Weka machine learning segmentation algorithm has the following steps:

#### Model training

– Define segmentation classes and select training data for each class from a training image.
– Compute selected image features for the training image.
– Extract image features and class for each training set voxel, generating a data table.
– Train classification model on this training data table using a selected Weka algorithm.

#### Model deployment

– For each image stack, compute the image features that are required for the trained segmentation model.
– Extract the image features for each voxel and apply the trained model to classify the voxel.
– Combine voxel classifications to produce a segmentation and (optionally) a probability map giving the estimated probability distribution over the classes for each voxel.

The role of the image features, using algorithms provided by the ImageJ platform and the ImageScience plugin (Meijering, 2015), resembles that of the earlier convolutional layers in the deep learning models sometimes used for semantic segmentation (Zinchuk & Grossenbacher-Zinchuk, 2020). However, using predefined image features radically reduces the cost of training in computational time and in the requirement for manually segmented training data, although the modeller must ensure that the selected features and scales capture sufficient spatial context for pixel classification. Full manual segmentation to produce training data may represent weeks of effort for a biologist, and represents a major limitation in the use of deep learning; since the generalisability of the model to new data is uncertain, potentially limiting the useful life of the model, the required effort is often impractical. The Trainable Weka approach uses a much smaller set of manually selected training data, although high quality results typically require assessment on a representative set of full images, and model iteration.

The Trainable Weka plugin uses the ImageJ interactive environment for both training and application of segmentation models in both 2D and 3D, which is highly effective for smaller image sets. Trained models may also be saved from the interactive environment, and either reloaded or deployed via non-interactive scripted processes. We encountered several bottlenecks when attempting to apply this software at scale: image features in 3D are expensive in computational time leading to long delays during the interactive process; available RAM limits the number of image features that can be used; software instability was encountered under heavy load, compounding the previous issues; training data selection was often difficult due to a lack of 3D context; and extending or revising existing training data selections, or combining training data for multiple stacks, is only possible using an ad-hoc process outside the interactive environment. We created our LLAMA visualiser, in the first instance, to provide clear 3D context during training data selections: see Supplementary Video 1. The other issues identified motivated our reengineered pipeline, in which the Trainable Weka plugin is used for the interactive selection of training data, but all computation is done non-interactively. Other key modifications are listed below:

- Image features are generated in a stand-alone process and cached to disk to minimise computational cost.
- **Selected image features may be approximated using a down-sampled image.** Features are calculated on a range of selected scales (the parameter *sigma*), and the computational cost increases with scale. But larger scales may be required to capture the required spatial context. We can effectively approximate larger scale features by down-sampling the original image, calculating the feature with appropriately reduced sigma, then up-sampling with interpolation. This greatly reduces computational cost. Importantly, the processes and code provided ensures that the features are calculated in a consistent way during training and deployment.
- **Training data selections are recorded in an ImageJ macro.** This process allows for editing and documentation of the selections and an easy way to resume or extend selections. It also enables the extraction of training data features to be handled by a separate non-interactive process with access to the macro file.
- **Full flexibility is allowed in feature selection.** In the plugin, each selected feature is used at each selected scale; the modified process allows any combination of feature and scale if desired.

### Object detection and quantification

Our novel object detection and tracking system involves an initial watershed split process (Beucher & Lantuéj, 1979) to separate touching objects, which should be calibrated to err towards over-separation to ensure that all necessary splits are implemented. This is followed by selective re-merging integrated with the tracking algorithm, allowing information sharing between time steps. In order to achieve scalability and high throughput, we took the approach of performing as much calculation as possible in parallel, on each stack in isolation, and then applying the tracking algorithm across all time steps as a separate step using summary object data. The computations performed in parallel for each stack include the expensive watershed split, as well as the calculation of a range of object metrics, which are aggregated appropriately when objects are merged during the tracking process.

The watershed algorithm selected was extended maxima watershed on the edge distance (Qin, Wang, Liu, & Yuan, 2013). Edge distance is the distance from each voxel in an object to the nearest external voxel; the idea is to use this metric to identify and remove narrow connections between more compact regions that are likely to represent separate objects. Classical watershed uses local maxima (equivalently minima) as seeds of expanding regions, but irregular object shapes may result in multiple local maxima in an object, leading to excessive splitting. The extended maxima approach solves this problem by specifying a minimum difference in the underlying metric (edge distance in this case) between adjacent regions. This minimum difference is the parameter “dynamic”. Given any two local maxima *m*_l_ and *m*_2_, they will be merged into the same region if there is a path between them where the minimum value is greater than min(*m*_l_, *m*_2_) − *dynamic*. The dynamic parameter is used to tune the algorithm, with smaller values leading to more splitting.

The watershed algorithm implementation (ExtendedMinimaWatershed) was provided by the MorphoLibJ plugin (Legland, Arganda-Carreras, & Andrey, 2016), working with 3D distance maps (edge distance transform) provided by the 3D ImageJ Suite (mcib3d-suite) plugin (Ollion, Cochennec, Loll, Escudé, & Boudier, 2013); 3D distance maps were also used to provide auxiliary statistical information on proximity between structures of different classes. The 3D ImageJ Suite plugin was additionally used for 3D hole filling, and to produce object meshes using the marching cubes algorithm (Lorensen & Cline, 1987) followed by mesh pruning. Skeleton representations of linear structures are produced using the plugins Skeletonize3D and AnalyzeSkeleton (Arganda-Carreras, Fernández-González, Muñoz-Barrutia, & Ortiz-De-Solorzano, 2010). Skeletons and meshes provide an efficient means of object visualisation and may also be used in statistical analysis.

A range of metrics are calculated for each object, including three that are used in the tracking algorithm: position, volume and the amount of contact with each adjacent object. This summary data is saved in tabular form for each stack, and forms the inputs for the tracking step described below.

### Tracking algorithm

In this section we provide a technical description of the tracking algorithm. Detailed instructions for applying the algorithm are included in the supplementary protocols, but the following text should also be consulted as a guide to selecting the key parameters used and understanding the output. We conclude with a brief guide to using the LLAMA visualiser to assess tracks and revise parameters.

The algorithm presented is designed to deal with data where adjacent structures often touch, the identity of individual structures is ambiguous, and methods used to delineate them can give inconsistent results between consecutive time steps. For example, it may be unclear at what point a dividing structure becomes two structures, or a disappearing structure completely recedes into the cell membrane. Biological variation between time steps or noise in the observational process means that a fixed algorithm applied to each time step may produce inconsistent segmentation results, leading to low quality tracking of structures over time, and the goal of the following is to correct these.

The approach is to pool information across time to give temporally stable and coherent representations of structures. We start with object data for individual time steps where an algorithm such as water-shedding has been used to separate touching structures, and the resulting objects have been reduced to a summary form; this has the advantage of allowing most computation to be done at the level of individual time steps, enabling parallel processing with relatively modest resource requirements. The splitting algorithm should be run under aggressive parameters to ensure that all true structures are split apart, at the cost of initially incorrectly subdividing many structures (if it is possible to completely avoid both under and over splitting, this tracking approach is not required and a simple matching algorithm is sufficient). We then selectively *merge* adjacent objects and *match* between successive time steps to form coherent structure tracks, seeking a balance between merging and matching in a sensible way and maintaining tracks over time.

The algorithm is customised with the following 6 parameters. Tracking is performed separately for each class of structure, so these parameters are specified for each class individually. See header information in the script “get_tracks.groovy” for details of how to set these and other parameters.

The general approach is to assign scores for merging adjacent objects at the same time step, based on the contact area, sizes and centre of mass distance between the objects, and combine with a distance metric for matching objects between consecutive time steps (with a threshold above which no match is made). We start by running a matching algorithm between consecutive time steps, forming a draft set of tracks (modified Hungarian algorithm (Kuhn, 1955) with distance threshold). Then we run an iterative process of track merging.

#### Quantification of object matches between adjacent time steps and merges at each time step

The distance metric used to match objects across time takes relative size into account as well as distance, with the relative importance controlled by the parameter *W_l_*. Given objects *a* and *b* with positions *p_a_*, *p_b_*, and volumes *ν_a_*, *ν_b_*, then the distance is defined as

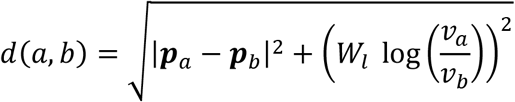

We require *d*(*a*, *b*) < *d_max_* for a match to be allowed. In order to help evaluate a set of tracks produced by matching and merging operations, we allocate a matching score

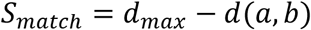

This score penalises poorer matches while rewarding longer tracks, since any match within the threshold gives a positive score.

The object merging score (defined below) is a metric designed to indicate whether two adjacent objects with the same class and time step are truly distinct, or if they should be merged and treated as a single structure. This score is a weighted average of two factors. The relative contact *R_e_* is an estimate of the contact area as a proportion of the surface area of the smaller object, which is approximated by the surface area of a sphere of the given volume (the lower bound of true surface area); both are in pixel units. The relative proximity *R_p_* is the inverse distance between the object centres with a correction factor based on the object volumes, since a given distance between centres of mass is indicative of a greater degree of separation for smaller objects. To get this correction factor, for each object we calculate the radius of the sphere with the same volume as the object, then we sum these 2 radii. This sum is then divided by the distance between the centre of masses to give *R_p_*. Given objects *a* and *b* with volume and position as above and contact area *C*(*a*, *b*), the formulae are

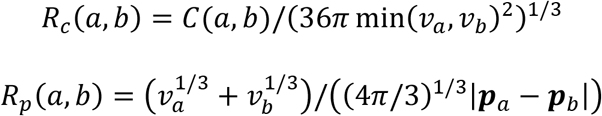

These factors are divided by reference values *Q_e_* and *Q_p_*, indicating the neutral level at which the score is not thought to be evidence for or against aggregation. The final merging score is a weighted sum of these factors, with weights based on the estimated usefulness of each.

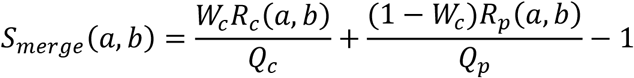

Final merge scores above 0 are taken as evidence for aggregation, but negative scores may still be consistent with aggregation when balanced by other score terms. The overall objective function is then a weighted sum of the scores of all merging and matching operations:

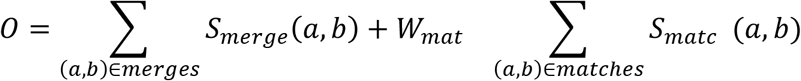

#### Tracking and iterative merging process

We seek to create a set of tracks by selectively merging objects at each time step and matching objects between consecutive times, in order to maximise the objective function above. This optimisation problem is computationally infeasible to solve in general, so we first solve the problem for matching only, then iteratively modify the result by merging tracks, allowing for matches to be added or removed in the process (splitting or joining tracks across time).

The objects are assembled into an initial set of tracks by matching objects at each pair of consecutive time steps. Matching distance is typically subject to a reasonably strict threshold (*d_max_*), such that we expect an object to be matched to at most one object in each temporal direction. But we account for the possibility that an object may have more than 1 possible match within this threshold, adapting the Hungarian algorithm to ensure that matches are unique (and hence tracks are simple / unbranched), and that total matching distance is minimised within these constraints. To do this, we first calculate the distance matrix between the 2 object sets. Then we replace all values greater than *d_max_* in the matrix with *d_max_*, and pad the smaller dimension of the distance matrix with dummy objects, with all associated distances set to *d_max_*. We then apply the Hungarian algorithm to this square matrix, and finally discard all distance *d_max_* matches and the dummy objects.

The initial set of tracks is then iteratively modified by track merging operations, in which two or sometimes more tracks, existing at the same or overlapping time periods, are merged together over some time interval [*t*_1_, *t*_2_]. Although there may be more than two tracks involved, exactly two must exist at each time between *t*_l_ and *t*_2_, and between 0 and 2 tracks may extend beyond the merged interval in each direction. The merge operations may involve one or more cuts to the tracks at the ends of the merged interval; if 2 tracks continue before or after the merged interval, at least one must be cut to avoid track branching. A prospective merge operation is scored by adding the merge scores at each time step and the change in matching score (the scores of all new matches made minus the scores of all discarded matches) weighted by *W_mat_*.

We begin the iterative merging process by considering every pair of tracks which are adjacent for at least one time point (touching objects). We find and score the optimal merge between the two tracks, by considering all possible merge intervals in the period where both tracks exist. For each possible merge interval we calculate whether continuing tracks should be included into the merged track or cut into separate tracks, in order to give the best match score adjustment, and this adjustment is included in the score for the interval. In the case where the merge score is positive but the overall score is negative after match score adjustment, we consider extending to further tracks (since this situation may be an artefact of a single point tracking failure). If the optimal merge interval continues to the end of the common time period, but one track continues beyond this time, then we look for tracks that are adjacent to the continuing track and start immediately after the merge period. If the matching at the end of the merge interval is improved by this new track, it is added to the hypothetical merge operation, and the merge interval is extended. This extension is continued in both directions while possible, adding new tracks as indicated, unless an overall positive merge score is achieved.

We then identify and execute the potential merge operation with the highest score, provided the score is positive. If possible, we extend the merged track by matching to existing tracks. Merge scores are then recalculated for the merged track, and any tracks formed by splits, and the highest merge score in this adjusted list is found. The process continues until no merges with positive score are available.

#### Assessing tracks and revising parameters using visualiser

The LLAMA visualiser may be used for assessing tracks, and in combination with the information above it may be used in setting or revising the tracking parameters: see Supplementary Video 2. A manual is included with the code repository, and the protocols contain a general guide for use in support of the analysis pipeline. Note that multiple track sets based on the same object data can be compared. In the data specification screen, select 2 or more object datasets with the same object folder but different track files, and provide labels to allow easy identification when the visualiser is running.

Useful tools include the “Selected” filter with specified track id and the “multiple times” option. Using “Object Coloring” you may see the objects as split by the watershed algorithm (“Object” option) versus the possibly merged structures produced by the tracking algorithm (“Node” or “Track” option; these are provided as separate options to support branched tracks if required).

A special feature is activated with the ‘c’ key in the visualiser. When showing tracks, the values of *R_p_*(*a*, *b*) and *R_e_*(*a*, *b*) are shown for each pair of touching nodes. Both values are multiplied by 100 and rounded to the nearest integer, then displayed as a pair *R_p_*(*a*, *b*) / *R_e_*(*a*, *b*) between the two nodes. By comparing the displayed values to your judgement of whether objects are best regarded as one structure or two, the appropriate reference values *Q_p_* and *Q_e_* can be selected.

## Supporting information

supplementary protocols

visualiser demo 1

visualiser demo 2

## Acknowledgements

LLSM imaging was performed using the resources of IMB Microscopy, which was partially funded by Australian Research Council Linkage Grant LE170100206, and the Australian Cancer Research Foundation funded Cancer Ultrastructure and Function Facility. Some of the LLSM datasets were acquired at AIC Janelia and we acknowledge the support of HHMI and the Gordon and Betty Moore Foundation. Most of the LLSM datasets were produced at IMB with additional technical support from 3i. We wish to acknowledge The University of Queensland's Research Computing Centre (RCC) for its support in this research in providing high performance computing facilities. Research funding was from the Australian Research Council Discovery Grant DP180101910 (NAH, JLS, AAW) and from the National Health and Medical Research Council Investigator Grant APP1176209 (JLS).

