## supplementary protocols for "LLAMA: a robust and scalable machine learning pipeline for analysis of cell surface projections in large scale 4D microscopy data"

### Overview

This document provides detailed instructions for the image analysis pipeline described in the Methods and Materials, including use of the visualiser. Necessary code is provided via Github:

- github.com/jameslefevre/4D-microscopy-pipeline
    - README file with installation instructions and other information, including description of some additional code that is not discussed here.
    - Compiled Fiji extension code Segmented_Image_Analysis.jar to be copied into Fiji jars folder (Java source code also provided).
    - A collection of parameterised groovy scripts used to control computations as described in the protocols below; documented in headers.
    - A template version of the file feature_model_table.txt, which contains information on segmentation models and the image features they use.
    - Template PBS job files for deployment of computations on HPC clusters (acting via groovy scripts above).
    - The file script_documentation.md containing detailed information on use of the PBS scripts.
- github.com/jameslefevre/visualiser-4D-microscopy-analysis
    - README file with installation instructions and other information.
    - Manual for visualiser app (visualiser_manual.md); provides more general and complete information than is provided here.
    - Visualiser source code in the Processing language.

The primary pipeline consists of the following protocols: (1) training a segmentation model; (2) deploying the segmentation model on a HPC system, allowing parallel processing of image stacks; (3) deploying the object splitting and analysis algorithm on HPC, designed to follow directly from the segmentation and retaining the parallel processing structure; (4) apply the tracking algorithm to the combined summary output from step 3, integrating information across time. The machine learning approach is designed for rapid training using a small training data selection, but high-quality segmentation results will often require an iterative process involving assessment of representative segmentations, followed by updating the model with additional training data. Protocol 5 is provided for this process. Protocol 6 provides instructions for using the provided visualiser to view outputs and support the main pipeline. Excluding Protocol 5, these correspond in order to the 7 system components listed in Table 1, with Protocol 3 including components 3, 4 and 5.

### Protocol 1: Train Weka segmentation model

The purpose of this protocol is to produce a Weka model file (Witten, Frank, Hall, & Pal, 2016), suitable for segmenting images from the target dataset into meaningful classes (in combination with the appropriate image features). This uses the Trainable Weka (Arganda-Carreras et al., 2017) plugin for ImageJ; see cited paper or the web page (Ignacio Arganda-Carreras) for an overview on using Trainable Weka.

If an earlier Weka segmentation model produced via this protocol exists, but is only partially suitable and requires modification, see Protocol 5 “Revising segmentation model”.

#### *Materials*

- Fiji software application (Schindelin et al., 2012), a distribution of ImageJ, with the provided file Segmented_Image_Analysis.jar copied into the “jars” folder, and the following plugins installed: Trainable Weka / Trainable_Segmentation, 3D_ImageJ_Suite / mcib3d-suite, MorphoLibJ, Skeletonize3D, AnalyzeSkeleton.
- Scripts generate_save_features.groovy, generate_training_data.groovy.
- Your version of feature_model_table.txt, or a copy of the template provided in the code repository.
- Image data to be used in training the segmentation model, consisting of one or more single-channel, 32-bit tiff stacks.

#### *Select training data*

1. Select and open image stack in Fiji.
2. Open new macro (Plugins>New>Macro) and save in the working directory, using a name associated with the current image stack. Additional information can be added at the top of the macro file in comments (start line with //). The file extension should be ijm.
3. Start the macro recorder (Plugins>Macros>Record; in the “record” option at the top left of the Recorder window, make sure “Macro” is selected.
4. Open the Trainable Weka 3D plugin (Plugins>Segmentation>Trainable Weka Segmentation 3D).
5. In the main Fiji window, click on the elliptical selection tool.
6. Returning to the Trainable Weka window, click settings and rename the two initial segmentation classes and click okay; then add and name any additional classes using the “Create new class” button. The first channel should be background.
7. Load the image stack in the visualiser (see Protocol 6), as an aid to correctly labelling regions (Optional).
8. Identify representative regions belonging to each class, make elliptical selections from each and add to appropriate class. *Aim for many selections from each class, in a full range of contexts, while being careful to avoid mislabelling. It is important to include background, sampling obvious background regions far from tissue as well as regions close to the tissue and any areas containing visual noise which are at risk of being mis-classified as tissue.*
9. Regularly cut and paste the text from the recorder window into the macro window (from step 2) and save. Errors may be corrected by editing the text, superfluous commands such as a selection not followed by an addition to training data may also be deleted, and comment annotations may be added if desired (start line with //).
10. Repeat steps 1-9 for each image stack used to produce training data, producing a macro file for each stack. Ensure that the classes at step 6 are consistent.

Fig. S1 shows the three windows used during the training data selection process. The data selection on a stack may be resumed after interruption by opening the image stack and corresponding macro, then running the macro and restarting the macro recorder. The macro can also be edited before running to revise the selections. This process also allows previous data selections to be extended when producing a new version of the segmentation model.

#### *Generate training dataset*

11. If starting a new segmentation model, add a column to the document feature_model_table.txt indicating the required features for the model. If working from the provided template version of this file, it may need to be substantially rewritten to reflect your choice of feature. The column header should be the unique name for the model being built. Any combination of the features provided by Trainable Weka and scale parameter (sigma) may be selected. In addition, a down-sampling factor may be specified for faster calculation at larger sigma (this gives an approximation, but this protocol ensures consistency between features in training and deployment). Note that the feature name acts as a unique identifier, while the following 5 columns define how the feature is calculated.
12. In Fiji, open the script generate_save_features.groovy and run in turn for each of the image stacks used to select training data (steps 1-9). You will be prompted to provide several parameters when you run; see the script header for parameter definitions and technical notes. The Feature_model_table path and ModelName must match the document edited in the previous step and the name of the column added. Leave cropbox blank if no cropping is required. Use a new folder for FeatureSavePath, and note the name. Set intensityScalingFactor to adjust intensity level to a selected benchmark level. This step generates the full set of selected features for the whole of the selected image stack, and saves to disk. *This step may take some time, however the saved features may be reused for modified training data selections.*
13. In Fiji, open the script generate_training_data.groovy and save to your working folder with a descriptive name indicating the origin and version of the training data (this serves as a record of the data source). Keep this file open for editing.
14. For each image stack that was used to select training data (steps 1-9), note the image file path, the path for the features generated in step 12, and the path for the macro used to record the selections (step 2,9) by adding entries to imageStackLocations, imageStackNames, featureFolders, and selection_macro_paths. Make sure to maintain a consistent order in these lists.
15. Set intensityScalingFactor for each image stack (set to 1 for no scaling). This can be used in place of the intensity scaling in step 12. It can also be used to augment the training data by replicating with rescaled intensity, to increase the robustness of the model to variations in intensity (since analysis should never depend on absolute fluorophore intensity levels). To do this, all the details for each image stack must be repeated for each scaling factor used.
16. Set the feature_model_table and modelName as for step 12, and classNames to match the names of the segmentation classes defined in step 6. Finally, give a path to save the training data, with file extension .arff, and run the macro.
17. Optionally, the training data can be generated in chunks, repeating steps 13-16 in each case. Feature selection and classNames must be identical. The resulting arff files must be combined by pasting the data from additional files (all text below the line @data) to the bottom of the original file. The text above this is the header information, which must be identical between all merged files and should not be

replicated. Manually editing these files may exceed the capacity of some text editors.

*Generate segmentation model*

18. Open the Weka Explorer app. *This is not the Trainable Weka plugin, but can be opened from it: click the Weka button on the lower left corner, then select Explorer from the popup window. Weka may also be installed and run separately.*
19. Open the arff file produced in step 16. If using a subset of the features in the data, remove those that are superfluous.
20. In Pre-process tab, choose filter supervised > instance > ClassBalancer (default parameter -num-intervals 10). Check that “class” is recognised as the Class variable, then click the Apply button
21. Select the “class” field to confirm it is now balanced. Optionally check class counts, compare/update training data counts in model.
22. Go to the Classify tab and select default random forest with 10-fold cross validation (other algorithms and parameters may be used, but this is a good default).
    Confirm class selected as target feature, and press Start.
23. Right click on trained model in the Result list and save the model with a name that can be linked to the training data, features, and algorithm used.
24. Right click again and save results buffer, and optionally open log and save that too; file names should be associated with the model file from the previous step. *Results may be compared here and different models may be produced with the same data (cross validated error rate is the best metric), but note that the training data is biased since it is selected manually, and the performance metrics give no guarantees about generalisability to the rest of the image data.*

Weka also contains tools to evaluate feature importance, which may help in reducing the number of image features used.

A

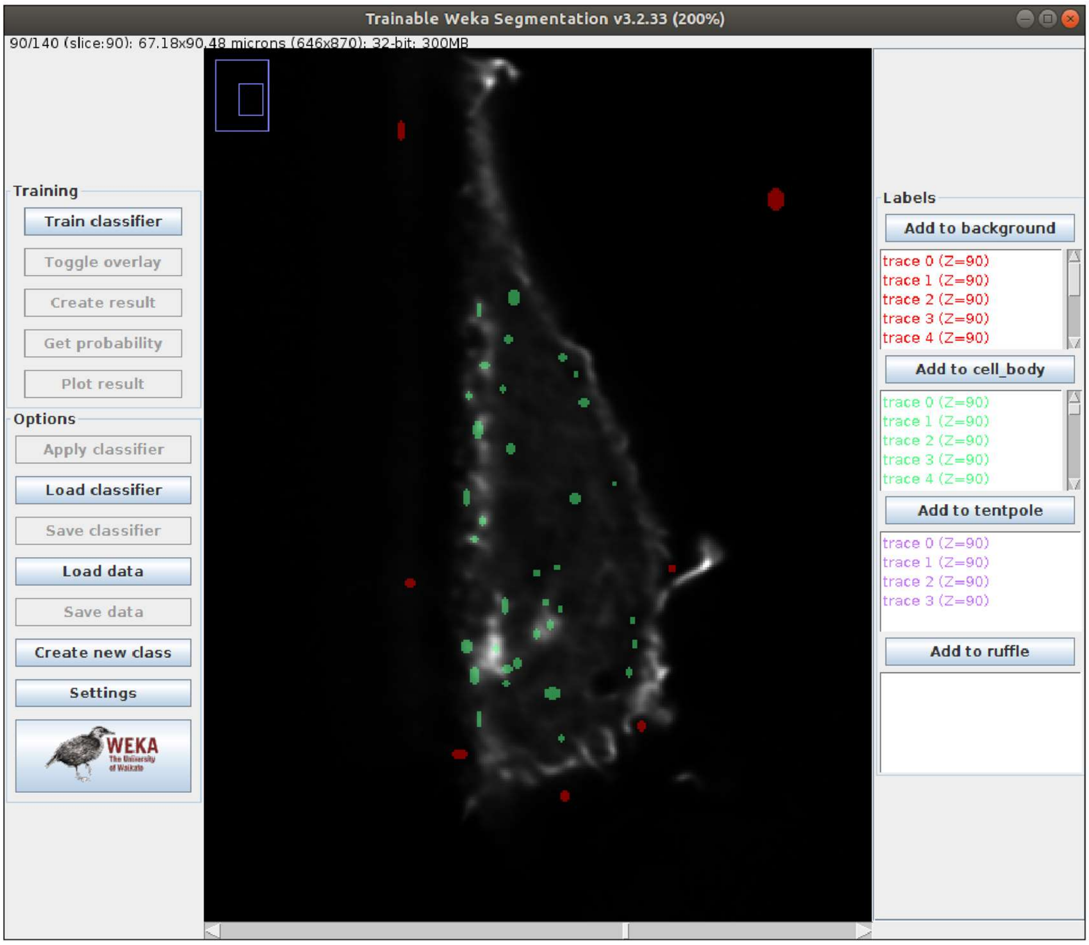

Trainable Weka Segmentation v3.2.33 (200%)
90/140 (slice:90); 67.18x90.48 microns (646x870); 32-bit; 300MB
Training
Train classifier
Toggle overlay
Create result
Get probability
Plot result
Options
Apply classifier
Load classifier
Save classifier
Load data
Save data
Create new class
Settings
WEKA
Labels
Add to background
trace 0 (Z=90)
trace 1 (Z=90)
trace 2 (Z=90)
trace 3 (Z=90)
trace 4 (Z=90)
Add to cell_body
trace 0 (Z=90)
trace 1 (Z=90)
trace 2 (Z=90)
trace 3 (Z=90)
trace 4 (Z=90)
Add to tentpole
trace 0 (Z=90)
trace 1 (Z=90)
trace 2 (Z=90)
trace 3 (Z=90)
Add to ruffle


B

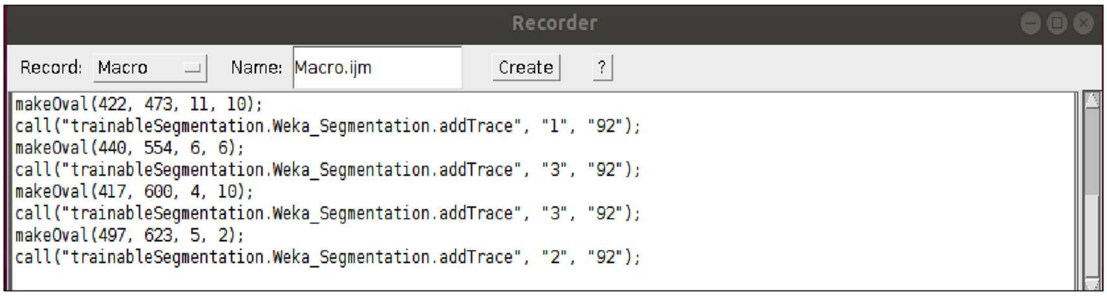

Recorder
Record: Macro
Name: Macro.ijm
Create
?
makeOval(422, 473, 11, 10);
call("trainableSegmentation.Weka_Segmentation.addTrace", "1", "92");
makeOval(440, 554, 6, 6);
call("trainableSegmentation.Weka_Segmentation.addTrace", "3", "92");
makeOval(417, 600, 4, 10);
call("trainableSegmentation.Weka_Segmentation.addTrace", "3", "92");
makeOval(497, 623, 5, 2);
call("trainableSegmentation.Weka_Segmentation.addTrace", "2", "92");


C

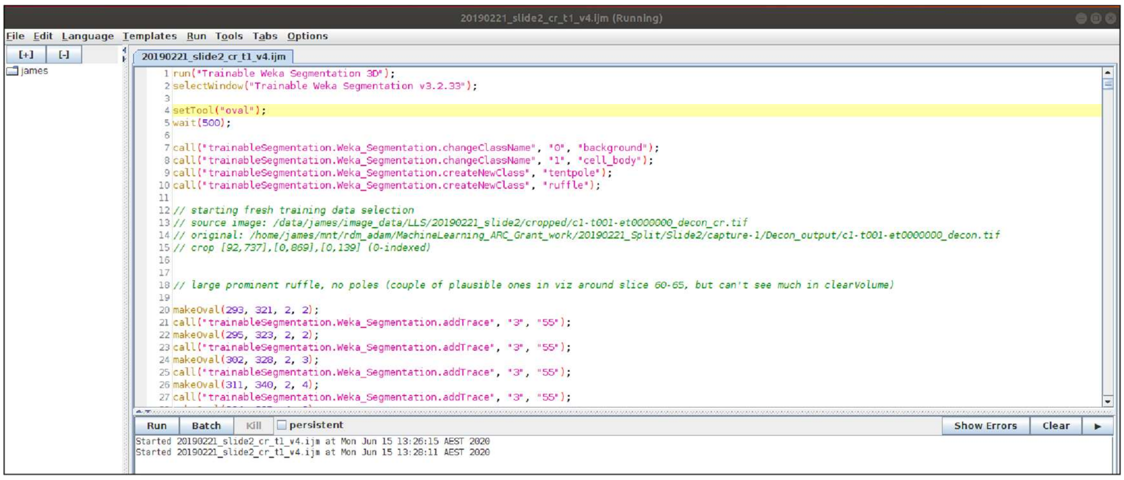

20190221_slide2_cr_t1_v4.ijm (Running)
File Edit Language Templates Run Tools Tabs Options
[+] [-]
james
20190221_slide2_cr_t1_v4.ijm
run("Trainable Weka Segmentation 3D");
selectWindow("Trainable Weka Segmentation v3.2.33");
setTool("oval");
wait(500);
call("trainableSegmentation.Weka_Segmentation.changeClassName", "0", "background");
call("trainableSegmentation.Weka_Segmentation.changeClassName", "1", "cell_body");
call("trainableSegmentation.Weka_Segmentation.createNewClass", "tentpole");
call("trainableSegmentation.Weka_Segmentation.createNewClass", "ruffle");
// starting fresh training data selection
// source image: /data/james/image_data/LLS/20190221_slide2/cropped/c1-t001-et0000000_decon_cr.tif
// original: /home/james/mnt/rdm_adam/MachineLearning_ARC_Grant_work/20190221_Split/Slide2/capture-1/Decon_output/c1-t001-et0000000_decon.tif
// crop [92,737],[0,869],[0,139] (0-indexed)
// large prominent ruffle, no poles (couple of plausible ones in viz around slice 60-65, but can't see much in clearVolume)
makeOval(293, 321, 2, 2);
call("trainableSegmentation.Weka_Segmentation.addTrace", "3", "55");
makeOval(295, 323, 2, 2);
call("trainableSegmentation.Weka_Segmentation.addTrace", "3", "55");
makeOval(302, 328, 2, 3);
call("trainableSegmentation.Weka_Segmentation.addTrace", "3", "55");
makeOval(311, 340, 2, 4);
call("trainableSegmentation.Weka_Segmentation.addTrace", "3", "55");
Run
Batch
Kill
persistent
Show Errors
Clear
Started 20190221_slide2_cr_t1_v4.ijm at Mon Jun 15 13:26:15 AEST 2020
Started 20190221_slide2_cr_t1_v4.ijm at Mon Jun 15 13:28:11 AEST 2020

**Figure S1: Selection of training data in ImageJ**
(A) Main window with image and Trainable Weka plugin open, showing segmentation classes and training data selections. (B) The macro recorder. (C) The macro which forms an annotated record of the selections made.

### Protocol 2: Semantic segmentation on cluster

This protocol describes the deployment of a Weka model produced according to Protocol 1 using an HPC (high performance computing) cluster. This allows image segmentation (see Fig. 2) to be performed on a large scale.

A segmentation model produced directly in the Trainable Weka plugin interface may be used in place of the model produced in the previous protocol, but the required features must be recorded in the file feature_model_table.txt as per this protocol.

#### *Materials*

- Fiji software set up as for Protocol 1.
- Scripts generate_save_features.groovy, apply_classifiers.groovy and segment_20190809_pre1_d18_intAdj_rep1ds1gd_rf (or another template job script), and the document script_documentation.
- Weka model file produced according to Protocol 1, and the feature_model_table.txt file containing information on the features used.
- Image data to be segmented. Each 4D dataset (image capture) consists of a directory containing a series of single-channel, 32-bit tiff stacks. The stack number / timestep must be included in each stack file name between 2 fixed text strings (the number may be padded with 0s on the left).
- Access to an HPC cluster using a PBS job submission system, with disk space for images to be processed. You should have basic knowledge of this system (how to access your home directory and copy files). Deployment on other HPC systems will need some programming/scripting knowledge.

#### *Apply segmentation model on cluster*

1. Copy the model file, the file feature_model_table.txt, and the scripts generate_save_features.groovy and apply_classifiers.groovy to your cluster home directory, and note the paths (from your cluster home directory root). Also copy over your Fiji folder (which includes the provided jar and required plugins) if it is on a local machine which is compatible with the cluster operating system (typically 64 bit Linux); otherwise a separate Fiji install is required for the cluster, set up as for Protocol 1. *Organise the copied files into subdirectories as convenient, but note the locations as the exact paths are required in the next step.*
2. For each capture (sequence of image stacks to segment), copy the file PBS array job file segment_20190809_pre1_d18_intAdj_rep1ds1gd_rf from the hpc_job_templates folder into your cluster home directory. Rename the file, and edit the PBS options, parameters and the paths to ImageJ-linux64, generate_save_features.groovy and apply_classifiers.groovy, to reflect your model, image capture location, file names and directory structure. *There are options to adjust image intensity to make it comparable with the training data (an initial scaling factor and an extra factor to apply again each time step to account for fade), crop the*

*images before segmentation, and to only segment files with a name that starts with a specified string. See script_documentation for details.*

3. Run the job script set up in step 2 as an array job (or sequence of array jobs) for the full range of stack numbers in the image capture. For example, the following generates jobs to segment stacks 100 to 150:
   qsub -q Short -J 100-150 [path to job script]
4. Monitor jobs and check for segmentations in the specified output directories when complete. For any failed jobs, analyse logs to identify errors or resources shortfalls and fix if possible, and rerun until all segmentations are complete. See your IT support for further help.

### Protocol 3: Perform object analysis on cluster

A semantic segmentation into tissue types does not in itself provide information on individual structures. This is because individual structures may be "stuck" together into a single object and need to be separated. We include code that extends the analysis pipeline with additional steps including a water-shedding algorithm to separate touching objects, the calculation of a range of metrics, and the optional generation of mesh and skeleton representations of objects (Fig. S2). The code extends ImageJ and plugin functionality and is designed to use as a scalable batch process. This protocol describes the deployment of this code on an HPC (high performance computing) cluster, allowing stacks to be processed in parallel for a scalable process.

Although designed as a continuation of the pipeline from Protocol 2, any segmentation may be used as input provided it satisfies the format requirements described below.

#### *Materials*

- Fiji software set up as for Protocol 1.
- Scripts split_object_analysis.groovy, get_meshes.groovy, get_skeletons.groovy, template job scripts objects_20190924_pre1, meshes_20190830_pos3_d19, skeletons_20190830_pos3_d19 (or previously adapted versions of them), and the document script_documentation.
- Segmented image data to be analysed. Each 4D dataset consists of a directory containing a series of single-channel, 8-bit colour tiff stacks. The stack number / timestep must be included in each stack file name between 2 fixed text strings (the number may be padded with 0s on the left).
- Optionally, the source imaging and segmentation probability maps. These are used to calculate object metrics and are not needed for the primary process. If used, they must be organised in separate folders with the same file names as the segmentations, and consist of single channel 32-bit tiff stacks and multi-channel 8-bit tiff stacks respectively.
- Access to a HPC cluster using a PBS job submission system, with disk space for images to be processed. You should have basic knowledge of this system (how to access your home directory and copy files). Deployment on other HPC systems will need some programming/scripting knowledge.

1. Copy the scripts split_object_analysis.groovy, get_meshes.groovy and get_skeletons.groovy to your HPC home directory, and note the paths.
2. For each segmented image capture to be processed, copy the PBS array job file objects_20190924_pre1 (or another version of this script) from the hpc_job_templates folder into your HPC home directory. Rename the file to reflect the source data including the model or segmentation version used, and open for editing.
3. Edit the PBS options according to the requirements of your HPC cluster and paths to match your file setup. Set parameters to select the required data and customise the analysis, and save. See script_documentation for details, in particular “primary object analysis”.
4. Run the job script set up in step 3 as an array job (or sequence of array jobs) to cover the full range of stack numbers in the image capture (or those that you wish to analyse). See script_documentation again for details. Job array index i will process stacks numberStacksPerJob*(i-1)+1 to numberStacksPerJob*i. For example, if numberStacksPerJob is 2, the following generates jobs to process stacks 1 to 20: qsub -q Short -J 1-10 [path to job script]
5. Monitor jobs and check the specified output folder when complete. For each processed stack, the output folder should have a subfolder with the name of the stack containing files objectStats.txt, objectAdjacencyTable.txt, classSummary.txt, and object_map.tif. If a class hierarchy is specified with more than 1 layer, there will also be additional files object_map2.tif etc. For any failed jobs, analyse logs to identify errors or resources shortfalls and fix if possible, and rerun until all stacks are complete. See your HPC/IT support for further help.
6. If object meshes are required, repeat steps 2-5 using template job script meshes_20190830_pos3_d19. Information on parameters is under “Generate object meshes” in script_documentation. Note that the required inputs are outputs from the primary object analysis (steps 2-5 above). The output files will be named objectMeshes.obj.
7. If object skeletons are required, repeat steps 2-5 using template job script skeletons_20190830_pos3_d19. Information on parameters is under “Generate object skeletons” in script_documentation. Note that the required inputs are outputs from the primary object analysis (steps 2-5 above), but this step is independent of step 6 (mesh generation). The output files will be named objectSkeletons.csv.
8. When all required meshes and skeletons are complete, the files object_map.tif, object_map2.tif etc may optionally be deleted to save disk space.

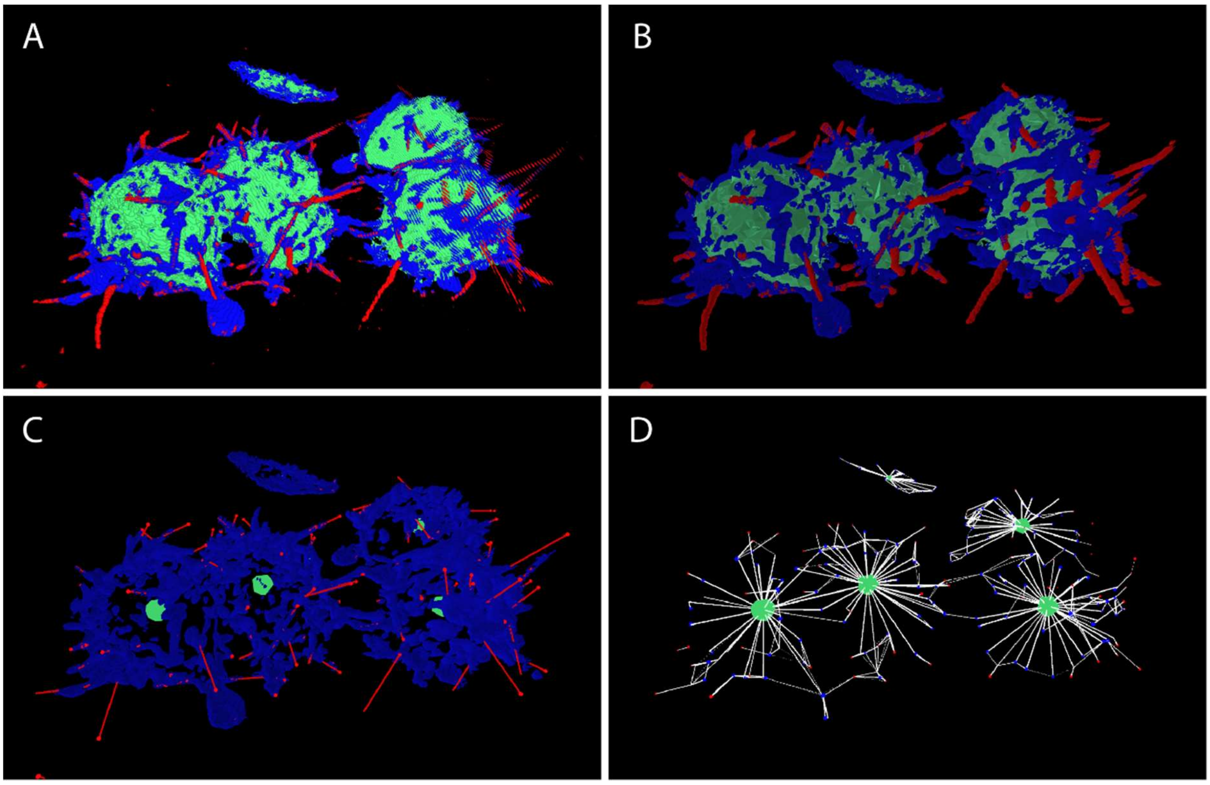


**Figure S2: Object analysis.**

The algorithm separates touching structures in the segmentation and produces a rich set of object information suitable for visualisation and statistical analysis. (A) A segmented image forms the starting point for the object analysis. (B) An object representation of the data in A, visualised with meshes; we see that the object representation successfully captures the important features of the segmentation. (C) An alternative object representation. Objects in the cell body class (green) are displayed using a sphere to show centre-of-mass position (sphere volume is reduced but indicative of true volume). Filipodia objects (red) are displayed using the skeleton representation, which is useful for linear structures, and provides length information for statistical analysis. The ruffle objects (blue) are displayed using object meshes, which are designed for visualisation of the true shape of structures. This provides a more computationally efficient representation of the segmentation which also allows for flexible display options such as filtering by size. Using the provided visualiser, the object representation allows extensive customisation of the information displayed. (D) Adjacency information can be used to establish relationships between structures. Lines are drawn between the centres of touching objects, with strength of line indicating the contact area.

### Protocol 4: Object tracking

This protocol builds on the object analysis in Protocol 3 in order to track structures over time (see Fig. 3). In Protocols 2 and 3, analysis is performed separately for each image stack (time step), to allow parallel computation and deployment to a cluster. Tracking requires information to be combined across time steps, so to avoid a computational bottleneck the tracking code works from a summary object representation for each time step.

Tracking is performed using a custom algorithm designed to give temporally consistent delineation of structures from a noisy process with ambiguous structure boundaries. This is done by selectively recombining split objects using information pooled between time steps. The algorithm and parameter details are given in Methods and Materials.

We provide a parameterised version of the tracking script, get_tracks_parameterised.groovy, to allow this process to be run through other code, including as a PBS job on a cluster (adapt PBS job scripts provided in Protocols 2 or 3).

#### *Materials*

- Fiji software set up as for Protocol 1.
- The script get_tracks.groovy
- The files objectStats.txt and objectAdjacencyTable.txt for each image stack to be included in the tracking, as produced by the object analysis protocol. A specification of these files is included in the get_tracks.groovy header. The data to be used must be arranged in a single folder containing a subfolder corresponding to each time step / stack. The name of each subfolder (typically the same as the image stack) must contain the stack number between two fixed strings (with optional zero padding to the left of the number), so that the number can be parsed without ambiguity.

1. In Fiji, open the script get_tracks.groovy and save to your working folder with a descriptive name indicating the data to be tracked, and if necessary information on parameter choices (this copy of the script can serve as a record of the derivation of the track set produced). Keep this file open for editing.
2. Edit the parameters in this script to select the data for analysis, the path for the output file, and the various parameters used in the tracking algorithm. *Parameters are documented in the script header, with additional explanation in Methods and Materials. The tracking algorithm is applied separately to each class, and the parameter values may vary between classes.*
3. Run the script and check the output file when complete. This should consist of a tab-separated text file holding a single table of "nodes". *A node consists of one or more of the objects in the input file, and is associated with a class, a track id and a timestep, as well as various ancillary data.*
4. Optionally, view the tracks in the visualiser tool provided. *This may also be used for quality assessment and revising parameters. See Protocol 6 for details.*
5. Perform statistical analysis of track data in a statistical environment such as R (the amount of data may be excessive for a spreadsheet). *The track data consists of a single table, with each row corresponding to a track node. Grouping variables include class, track id and timestep, and additional information is included depending on the parameters used in the object analysis computation.*

### Protocol 5: Revising segmentation model

While Protocol 1 gives complete instructions for training a segmentation model that can be deployed using Protocol 2, optimal segmentation results will generally require an iterative process with at least one cycle of evaluating and improving the model, typically by selecting additional training data in locations where the model underperforms.

Trainable Weka allows models to be developed rapidly using small amounts of selected training data, but this training data may not be completely representative of the full range of imaging to be processed, due to both limited quantity and the biased selection process.

Thus, even a model which performs well on the selected training data may not generalise well to the complete dataset.

This protocol describes this evaluation and revision process. It is also applicable where an existing model needs to be tweaked in order to perform well on new data with slightly different characteristics.

1. Produce a segmentation model according to Protocol 1.
2. Segment your data or a representative sample of it following Protocol 2.
3. Assess the segmentation in the visualiser (see Protocol 6), or elsewhere.
4. If the segmentation needs improvement, identify image stacks and specific location where mis-segmentations occur. Most importantly, identify examples of any widespread class of error.
5. Repeat Protocol 1, with the following modifications: (1) if selecting additional training data from a stack which has already been used for training, at step 2 load the existing macro (optionally, save under a new name), remove the code for any previous selections if you consider them to be in error, then run the macro, before continuing with step 3, 5 then from step 7. (2) Repeat step 11 only if you wish to change the selection of image features used. (3) Repeat step 12 only for image stacks which were not previously used for training data, unless you are adding new image features to the model, in which case repeat for all image stacks used for training. (4) For steps 13-16, update and run your existing script. *Training data selections should generally focus on misclassified voxels; the visualiser provides tools for efficiently identifying them.*
6. Return to step 3.

### Protocol 6: Use of visualiser

This protocol describes the use of the visualiser tool (Fig. S3), which is provided to support other processes described here. The visualiser is used in training data selection and the evaluation of segmentations and object representations, as well as visualisation of outputs. The 3D/4D visualiser is designed to facilitate comparisons between 2D and 3D views, data at different time points, and between raw image data, image segmentations, and object representations (potentially including alternative segmentations of the same data and alternative object representations based on the same segmentations). The various object representations provided by Protocol 3 can be viewed selectively, including showing object tracking across time.

There are three types of image data that can be displayed: the original imaging (single channel), class segmentations, and the probability maps associated with the segmentations (which shows the estimated probability that a given voxel is in each of the classes, rather than just giving the class which is considered most likely).

#### *Materials*

- Visualiser application
- Visualiser manual (included with visualiser). Includes installation and running instructions.

- Image and object data to visualise copied locally or on a mounted drive, in the format required / produced by the segmentation, object analysis and tracking algorithms; see manual for full specification.

#### *Data preparation*

1. This step is required in order to visualise any image data (does not apply to object representations). Open the script “export_to_vis.groovy” in Fiji and run once for each image capture to be visualised. Specify the source folders for the three image types (original, segmentations and probability maps) and for the corresponding destination folders, and set other parameters. *These parameters are documented at the start of the script. Step 2 should be considered when setting the destination folders. Note: use forward slash character (/) in paths; these will be translated to backslashes on Windows systems.*
2. Ensure that the image files produced in step 1 and any object files to be visualised are arranged according to the specification in the visualiser manual (“data specification” section). *Not all image types need to be present, and as the pipeline progresses additional output types can be added, following these steps.*

#### *Using visualiser*

3. Run the visualiser and specify data to load (see “Data selection” section in manual). Note that selected data must fit into RAM on your machine; image data is the most expensive, followed by meshes, while other image data (size, position, track, adjacency information and skeletons) uses very little memory. See the “User interface” section of the manual for help on viewing the data that you have loaded.
4. When selecting training data for a new model, display the original imaging and alternate between 3D and slice view in order to select training data in 3D context (the controls for this are at lower left of interface). Slice view allows direct comparison with the Trainable Weka interface where training data is selected. If segmentations have been generated and loaded into the visualiser, they can be compared to the source images in 2 and 3 dimensions to evaluate the segmentations, and additional training data selected as above to correct identified errors in a new model version. There is also a capacity to load and compare multiple versions of the segmentation.
5. After producing object representations corresponding to your segmentations (see Protocol 3 and 4), these can be loaded into the visualiser and compared to the segmentations. As well as providing a visualisation of results, this allows the quality of the object analysis to be assessed, and parameters to be revised if necessary. Multiple versions of the object analysis corresponding to the same segmentation can be loaded and compared.

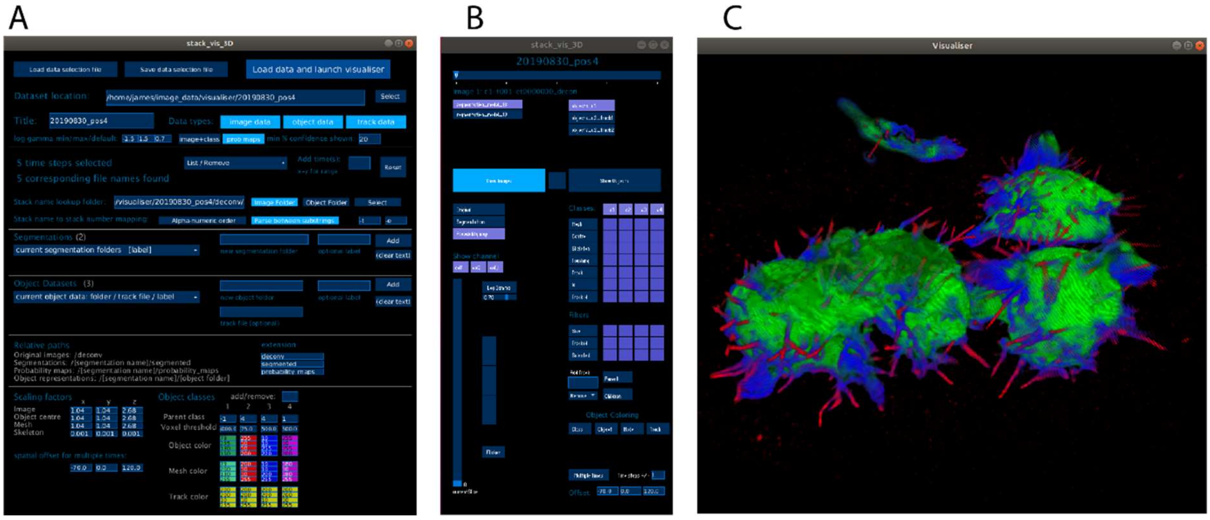

**Figure S3: Visualiser**

Interface for the visualiser tool. (A) Data selection window that is shown when the application starts. The selections control what image and object data is loaded into memory, and also provide some scale and colour customisation. When the launch button is pressed (top right), a temporary progress screen is shown during data loading, then the visualiser is launched and panels B and C are shown. (B) The control window that is displayed after launch alongside the visualiser window. Gives fine grained control of data to be displayed. (C) The visualiser window, displaying the data selected in the control window. Mouse controls allow zooming, panning and rotation.

### Critical Parameters

#### Intensity correction

Fluorophore intensity should be normalised between all analysed and training images. No specific protocol is included for this purpose, but the intensityScalingFactor parameter is provided in Protocols 1 and 2 (see script_documentation for details). In addition, training data may be replicated with scaled intensity to provide a model which is more robust against intensity variation: see Protocol 1, step 15. Normalisation between captures ideally uses a benchmarking strategy, since the mean intensity of an image will be biased by the amount and type of tissue included; in the example provided we used cytoplasm intensity, from the region select and measure tool in ImageJ (if the measured cytoplasm intensity is $I_1$ versus $I_0$ in the selected reference image, we use intensityScalingFactor=$I_0/I_1$ for the image). Intensity decay within a capture may be modelled using average intensity, provided there is no large movement of tissue in or out of the imaging region. Note that excess fluorophore degradation may lead to a changing pattern of illumination or loss of image quality that cannot be corrected by normalisation; we required that intensity remained above 50% of the initial value at a minimum.

#### Segmentation

As with other machine learning algorithms, the semantic segmentation models produced can only be relied upon for data that is sufficiently similar to the training data. Although we have found relatively good generalisation behaviour using random forest with a large range of features, no guarantees can be given. Changes to image characteristics such as the pattern of noise can lead to model failure, and care must be taken to ensure that training data is representative of the full range of variation in the image data. Protocol 5 outlines an iterative process to help ensure this is the case; the segmentation model is assessed using the provided visualiser, and revised using additional training data.

The image features used are calculated at multiple specified scales, and these should reflect the type of information required to classify voxels. In particular, the largest sigma parameter used represents the maximum extent of the context used in classification: if you consider the sphere of this radius around a given voxel, no information from outside this sphere can influence the class assigned to the voxel. Large sigma values can be computationally expensive, so we have included a mechanism to approximate features with selectively down-sampled imaging, avoiding the necessity of a general down-sampling (which is also an option). See Protocol 1, step 11.

### Training model using Protocol 1 versus standard Trainable Weka interactive process

Protocol 1 describes a modified version of the Trainable Weka model training process which is performed on a local machine and retains use of the Trainable Weka interface for training data selection. It is possible to train a model using the standard, fully interactive Trainable Weka process, but this protocol and associated code provides a modified process that is more scalable and extensible. The most important modifications are listed here as a guide to when this protocol may be required. Note that if you use a model produced using the standard fully interactive Trainable Weka process, it is still necessary to record the features used in the file feature_model_table.txt as per Protocol 1 in order to use Protocol 2 for deployment.

- **Image features are generated in a stand-alone process and cached to disk to minimise computational cost.** Training a model requires computational of the full image feature stack of each image used. This can lead to high memory use, since each feature requires as much space as the original image, and may be computationally intensive when using 3D features.
- **Selected image features may be approximated using a down-sampled image.** Features are calculated on a range of selected scales (the parameter *sigma*), and the computational cost increases with scale. But larger scales may be required to capture the required spatial context. We can effectively approximate larger scale features by down-sampling the original image, calculating the feature with appropriately reduced sigma, then up-sampling with interpolation. This greatly reduces computational cost. Importantly, the processes and code provided ensures that the features are calculated in a consistent way during deployment.
- **Training data selections are recorded in an ImageJ macro, allowing editing and documentation of the selections and an easy way to resume or extend selections.**
- **Full flexibility is allowed in feature selection.** In the plugin, each selected feature is used at each selected scale; the modified process allows any combination of feature and scale if desired.

### Object and track data

The algorithms provided to identify and track structures are designed to work robustly, without requiring editing of individual objects or tracks. However, a range of parameters need to be specified for each segmentation class, and the optimal values will depend on the characteristics of typical structures (mostly size and shape). Trial and error is likely to be required to find these optimal values. One function of the visualiser is to assess the object and track outputs to assist in this revision process. The most important parameters are the watershed split parameter “dynamic” (see script_documentation, primary object analysis section; smaller values of “dynamic” will result in more aggressive splitting of touching objects) and the tracking parameters explained in detail in Methods and Materials.

### Time Considerations

There are 2 main time considerations which are estimated separately: the time required for a person to carry out the steps (user time) and the time required for computation (which typically takes place as a background task). Computational times are based on a sample 12 million voxel image (225x517x104), which contains one large cell at 104x104x268nm resolution, which was segmented using 111 3D image features.

#### Model training

The estimated minimum user time for Protocol 1 is approximately 2 hours, but significantly longer times can be expected depending on the amount of training data selected and the difficulty of correctly labelling classes. Note that training data must be representative of the full range of data to be segmented, so the difficulty will depend on the size and complexity of the image data set, including variability in image properties and the intrinsic difficulty of separating segmentation classes. Refining the process using the iterative process described in Protocol 5 additionally requires use of Protocols 2 and 6 for generating and assessing draft segmentations, and for a difficult segmentation problem this may require multiple days of work in total. Testing of alternative models or model parameters in Weka can be performed within a few minutes to an hour, but testing generalisability to a larger set of images and comparing to other models is likely to take several hours.

In terms of computation time, generation of the 111 image features for the 12 million voxel sample image took approximately 8 minutes with a single threaded CPU process. This increases in proportion to the number and size of images used for training data. The computational time for training the random forest model was under 1 minute.

#### Segmentation

Given familiarity with the Linux command line and the HPC facility to be used, minimum user time for Protocol 2 should be approximately 15 minutes. This is to set up the segmentation for one image capture (sequence of image stacks), and it does not include any time taken to calculate intensity adjustment factors or cropping regions, which is out of scope for the protocol. This time also does not include copying image files to the HPC system, or the initial setup of the HPC environment, which will depend on available transfer speed. The time required may also increase if a significant number of jobs fail. This can result from inadequate resource allocation in the job script, or computational issues such as network interruptions.

Computational time for the 12 million voxel sample image was approximately 12 minutes on a single threaded HPC job, including 8.5 minutes to generate 111 image features and 3.5 minutes to run the model to classify each voxel.

#### Object analysis

Given familiarity with the Linux command line and the HPC facility to be used, minimum user time for Protocol 3 should be approximately 30 minutes. This includes the generation of object meshes and skeletons as well as the base object analysis, requiring 3 job scripts to be customised and run. Input files should be in place following Protocol 2. As for segmentation, this time does not include initial environment setup, and failed jobs will require additional effort to monitor results and re-run HPC jobs where required.

Computational time for the 12 million voxel sample image on a single threaded HPC job was approximately 1.5 minutes for the primary object analysis, 2 minutes for object meshes, and 8 seconds for the skeletonization (performed on the sparse filipodia class only).

#### Tracking

The generation of the tracks for a dataset should take only about 15 minutes user time, mostly setting paths and parameters. Computational time varies considerably depending on complexity, with some of our samples requiring over 1 hour (single thread background process on a local machine). The size threshold for including objects in the algorithm can be used to control complexity to some degree. Selecting the optimal values for the main tracking parameters may require trial and error and comparisons between different versions in the visualiser. Depending on the difficulty of the tracking problem this may take hours or days, although the same optimised settings should be effective for multiple captures if the data is similar in nature.
